# The *Drosophila* ZAD zinc finger protein Mulberry shapes the organization of the regulatory genome in the early embryo

**DOI:** 10.64898/2026.02.03.703437

**Authors:** Yusuke Umemura, Takashi Fukaya

## Abstract

Long-range regulatory interactions play a fundamentally important role in the control of gene activity during animal development, yet the underlying mechanisms remain largely unclear. Here, we identified a zinc finger-associated domain (ZAD)-C2H2 zinc finger protein, CG31365/Mulberry, as a looping factor that mediates long-range tethering activity in the early *Drosophila* embryo. Evidence is provided that Mulberry is specifically recruited to a subset of loop anchors and topological boundaries at key developmental loci to shape genome organization and gene activity. Super-resolution imaging analysis revealed that Mulberry forms nuclear condensates that associate with its target loci through the structured N-terminal ZAD domain. Micro-C analysis further demonstrated that the formation of loops and boundaries is lost in the condensation-deficient Mulberry mutant in a locus-specific manner. We propose that Mulberry acts as a condensation-dependent structural regulator of genome topology, organizing “multi-way regulatory hubs” that mediate long-range gene activation during early embryogenesis.

## Introduction

Long-range regulatory interactions play a critical role in controlling gene activity in time and space during animal development. In recent years, it has become clear that the organization of these interactions relies on two distinct classes of DNA elements: topological boundaries (or insulators) and tethering elements.^1–3^ In mammals, topologically associating domains (TADs) are thought to be shaped by the interplay between loop-extruding cohesin and boundary-associated CTCF,^4–10^ whereas in *Drosophila*, arthropod-specific zinc-finger proteins such as Su(Hw) and CP190 also contribute to 3D genome organization.^11–13^ In addition to these well-characterized factors, nuclear proteins such as Yin Yang 1 (YY1), LIM domain binding protein 1 (Ldb1), and POZ/BTB and AT hook containing zinc finger 1 (PATZ1) have recently emerged as key regulators that facilitate long-range loop formation in mammals.^14–27^ A common molecular feature of this class of looping factors is their ability to self-associate. For example, Ldb1-mediated looping at the *β-globin* locus requires a self-association domain at its N-terminus,^23^ and direct tethering of the Ldb1 self-association domain is sufficient to reactivate developmentally silenced genes by forming new loops in a cohesin-independent manner.^16–18,20–22^ Similarly, in *Drosophila*, the zinc finger DNA-binding protein GAGA-associated factor (GAF) mediates long-range tether-tether interactions through its self-associating POZ/BTB domain in developing imaginal discs and brains.^28,29^ More recently, another POZ/BTB-containing protein, Lola-I, was reported to promote the formation of long-range regulatory loops in the *Drosophila* brain.^30^ These loops have also been suggested to foster long-range co-regulation or cross-regulation of functionally related paralogous genes in developing embryos,^2,31^ underscoring the biological significance of such interactions. However, the relative contribution of each factor still remains uncertain*^e.g.^*^,32^ and the identification of new regulators represents one of the major challenges in the field.*^e.g.^*^,14,15,30,33,34^

As these examples illustrate, even in the relatively compact *Drosophila* genome, it is becoming clear that classically studied insulator proteins such as CTCF, Su(Hw), and CP190 cannot fully account for the complexity of regulatory genome organization.^35–38^ Although POZ/BTB-containing GAF and Lola-I possess looping activity, they appear to be responsible for only a small subset of loops during *Drosophila* development.^28,30^ For example, only ∼6% (12/186) of loops are selectively lost in POZ/BTB-deficient GAF mutant.^28^ A recent study identified the MADF transcription factor Vostok as an additional looping factor capable of mediating long-range tether-tether interactions in the *Drosophila* brain,^34^ yet similar to GAF, only ∼7% (47/645) of loops are selectively lost upon CRISPR-mediated knockout of *Vostok*. Locus-specific loss of topological boundaries is also observed upon depletion of well-studied insulator proteins such as CTCF, BEAF-32, and CP190 in developing embryos.^36^ Collectively, these findings are consistent with the idea that the regulatory genome is organized through the cooperative action of multiple locus-specific looping factors. Accordingly, identification and characterization of new regulators of long-range looping interactions is essential toward a full understanding of the basic principles of 3D genome organization.

Zinc finger-associated domain (ZAD)-containing C2H2 zinc finger proteins (ZAD-ZnFs) represent the most abundant class of transcription factors in insects including *Drosophila*.^39,40^ These proteins consist of an N-terminal ZAD and C-terminal arrays of C2H2 zinc finger domains, separated by a flexible linker region. Intriguingly, similar to the dimerizing POZ/BTB domain, biochemical and X-ray crystallographic studies have shown that the N-terminal ZAD possesses self-association activity,^41–44^ raising the possibility that ZAD-ZnFs may also function as locus-specific looping factors in *Drosophila*.

In this study, a comprehensive survey of stage-specific RNA-seq data identified *CG31365* as one of the most highly expressed ZAD-ZnF genes in early *Drosophila* embryos. Using a combination of genetic analyses, ChIP-seq and Micro-C assays, and super-resolution live-imaging, we provide evidence that CG31365, termed as Mulberry, functions as a condensation-dependent looping factor that is critical for 3D genome organization and gene regulation in the developing embryo.

## Results

### Identification of *CG31365* as a ZAD-ZnF gene with a key biological function

To explore previously uncharacterized ZAD-ZnF genes with key biological functions in early *Drosophila* embryos, we surveyed the expression levels of all ZAD-ZnF genes in nuclear cycle (nc) 14 embryos using stage-specific RNA-seq data.^36^ This analysis led to the identification of a poorly characterized *CG31365* as one of the three most highly expressed ZAD-ZnF genes in early embryos (Figure 1A). This finding attracted our particular attention because the two other abundant genes, *hangover* (*hang*) and *Motif 1 Binding Protein* (*M1BP*), encode DNA-binding proteins with essential functions in transcriptional regulation and genome organization.^45–50^ To date, no biological or molecular functions have been reported for *CG31365* despite its remarkably high level of gene expression. Like other ZAD-ZnFs in *Drosophila*,*^e.g.^*, ^51–57^ the CG31365 protein consists of an N-terminal ZAD, followed by a flexible linker, and a C-terminal array of C2H2-type zinc finger domains (Figure 1B). To investigate the physiological role of CG31365, we employed CRISPR/Cas9-mediated genome-editing to remove the entire transcription unit of *CG31365* (Figure 1C; *CG31365[Δ1]* denotes the resulting full-locus deletion allele). The *CG31365[Δ1]* mutant was found to be homozygous viable, but embryos from homozygous mutant adults exhibited a clear reduction in hatching rates (Figure 1D). Live-imaging analysis of mutant embryos further revealed developmental defects, including asynchronous mitosis and disorganized nuclei (Figure 1E), reminiscent of the phenotype observed in GAF-depleted embryos.^58^ Overall, our data are consistent with the idea that CG31365 exerts an important biological function during early embryogenesis.

**Figure 1.**
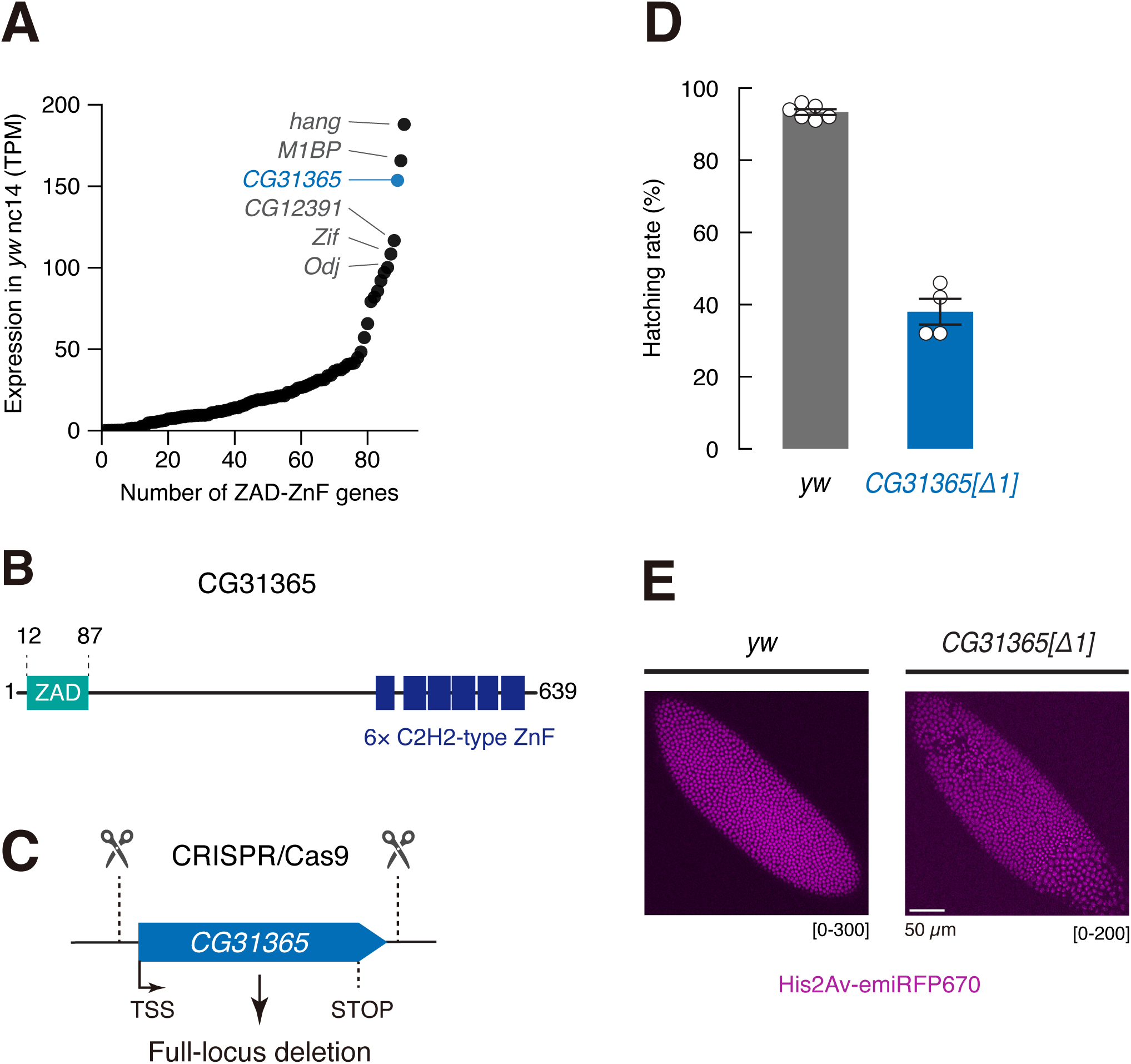
CG31365 is a ZAD-ZnF with key biological function. (A) Expression levels of ZAD-ZnF genes. Publicly available RNA-seq data in nc14 embryos^36^ (E-MTAB-11978) was used for the analysis. The expression levels were normalized to TPM (transcripts per million). (B) Schematic of the full-length CG31365 protein. (C) Schematic representation of CRISPR/Cas9-mediated full-locus deletion *CG31365[Δ1]* allele. (D) Measurement of the embryo hatching rate. Embryos from *yw* adults or homozygous *CG31365[Δ1]* mutant adults were used for the analysis. Bars represent the mean, and error bars represent the standard error of the mean. n = 6 (600 embryos) for *yw*; n = 4 (400 embryos) for *CG31365[Δ1]*. (E) Representative confocal images of His2Av-emiRFP670 in embryos with a genetic background of *yw* or homozygous *CG31365[Δ1]* mutant. Maximum intensity projected images of His2Av-emiRFP670 are shown.

### CG31365 forms nuclear condensates through its N-terminal ZAD domain

To elucidate the molecular properties of CG31365 in early embryos, we used CRISPR/Cas9-mediated genome-editing to insert a GFP-3×FLAG tag at the C-terminus of endogenous CG31365 (Figure 2A).^57^ Airyscan super-resolution live-imaging revealed that endogenous CG31365 forms bright nuclear foci during both interphase and mitosis (Figure 2B and Video S1). To characterize the molecular dynamics of CG31365 within these foci, we performed Fluorescence Recovery After Photobleaching (FRAP) in interphase nuclei. The GFP signal rapidly recovered after photobleaching (∼100 sec) (Figure 2C), indicating that CG31365 foci, or condensates, represent dynamic subnuclear assemblies. We then sought to determine which region of CG31365 is responsible for its condensation activity. Previous biochemical and X-ray crystallographic studies reported that the N-terminal ZAD possesses intrinsic self-association activity through the stacking of hydrophobic surfaces.^43,44^ Consistent with these findings, AlphaFold3 predicted that the N-terminal ZAD of CG31365 forms homodimers with high confidence (Figure 2D). To test the role of the ZAD domain in condensation, we generated transgenic strains expressing either full-length (WT) or ZAD-deficient (ΔZAD) CG31365 fused to GFP-3×FLAG under the control of the native *CG31365* promoter. Live-imaging analysis revealed that condensation activity is specifically lost in the absence of the N-terminal ZAD, resulting in uniform nuclear distribution of CG31365 (Figure 2E). Molecular phylogenetic analysis suggested that *CG31365* belongs to an ancient class of ZAD-ZnF genes conserved throughout 40 million years of *Drosophila* evolution (Figure S1A). Intriguingly, AlphaFold3 predictions suggested that the self-association activity of the N-terminal ZAD is conserved not only across *Drosophila* species but also in other insects such as mosquitoes (Figure S1B). Thus, it is tempting to speculate that CG31365 condensates play an active role in exerting its function across divergent insect species.

**Figure 2.**
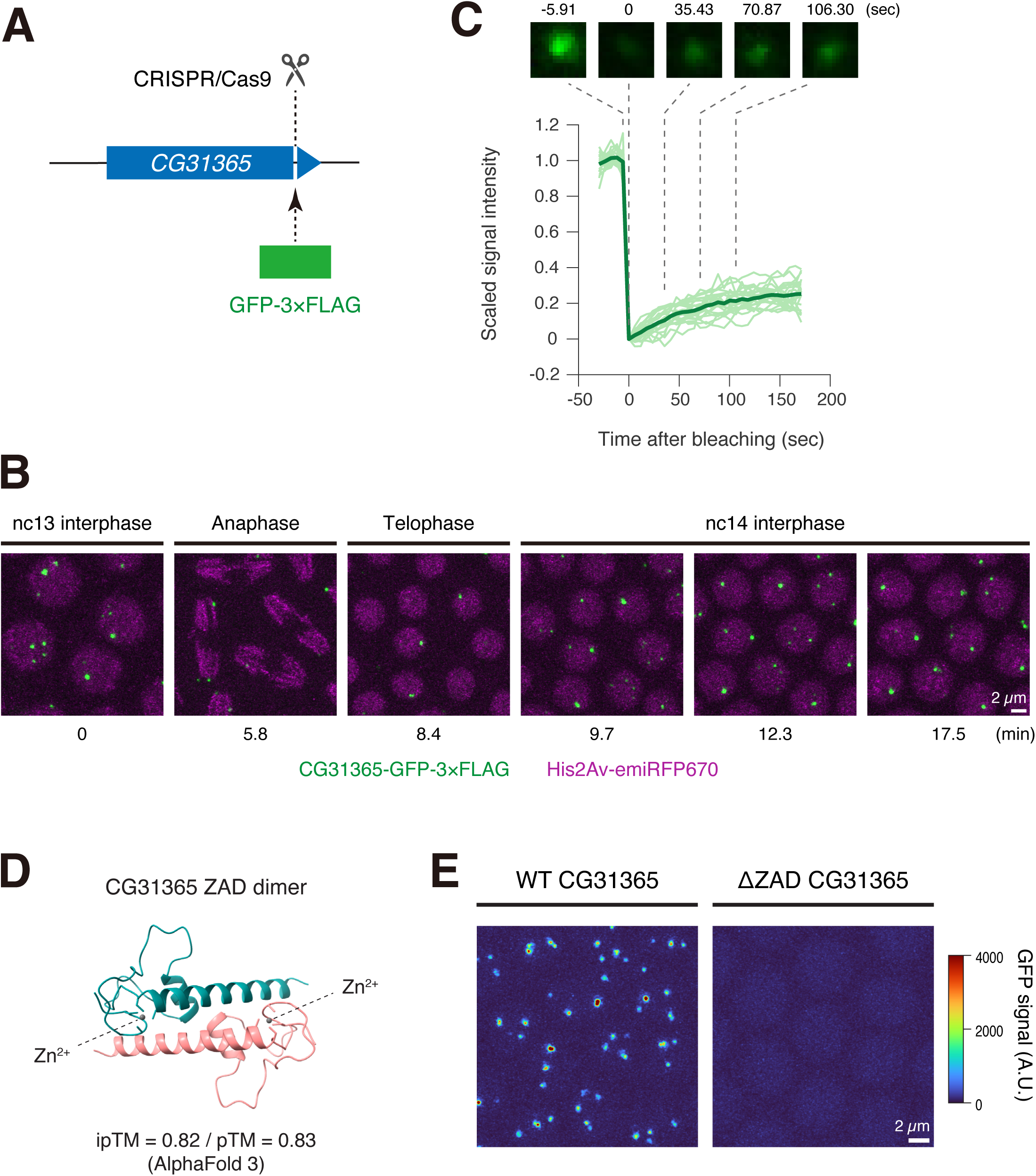
CG31365 forms dynamic nuclear condensates through ZAD domain. (A) Schematic representation of CRISPR/Cas9-mediated GFP-3×FLAG tagging for CG31365 protein. (B) Representative Airyscan time-lapse sequence of CG31365 localization pattern. Maximum projected images are shown. CG31365-GFP-3×FLAG and His2Av- emiRFP670 signals are shown in green and magenta, respectively. (C) FRAP analysis for CG31365 condensates. (top) Representative time-lapse sequence of a photobleached CG31365 condensate. Maximum projected, median-filtered images of CG31365-GFP-3×FLAG are shown. Images were cropped around the condensate. (bottom) Fluorescence recovery curve of GFP signal. Thin and thick lines represent signal profiles from each replicate and an averaged signal profile, respectively. 22 condensates from 13 embryos were analyzed. (D) AlphaFold3^77^ prediction of the CG31365 ZAD dimer. Protein structure was visualized using UCSF ChimeraX.^78^ pTM and ipTM values are used as proxies for the accuracy of predicting the overall protein folding and the relative positioning of the monomers within the dimer, respectively (hereinafter the same). (E) Representative Airyscan imaging of GFP-3×FLAG-tagged WT and ΔZAD CG31365 expressed under the control of the native *CG31365* promoter. Images were taken at ∼700 sec after entry into nc14. Embryos expressing transgenes in *yw* background were used for the analysis. Maximum intensity projected images are shown. See also Figure S1.

### CG31365 and CP190 co-occupy a subset of topological boundaries

Motivated by our recent finding that many ZAD-ZnFs are recruited to boundaries of topologically associating domains (TADs),^57^ we examined whether CG31365 similarly associates with topological boundaries in early *Drosophila* embryos. Notably, yeast two-hybrid and *in vitro* GST pull-down assays have suggested that CG31365 can directly interact with the insulator protein CP190.^59^ To investigate the genome-wide binding profiles of endogenous CG31365 and CP190 in early embryos, we performed ChIP-seq using a well-established anti-FLAG M2 antibody,^60^ targeting GFP-3×FLAG-tagged endogenous CG31365 and CP190 individually (Figure 2A and S2A).^57^ Our CP190 ChIP-seq dataset showed strong agreement with previously reported profiles generated using an antibody against endogenous CP190 (Figure S2B), validating our approach. Importantly, comparative analysis of the CG31365 and CP190 ChIP-seq profiles together with our Micro-C contact maps in nc14 embryos revealed a high degree of CG31365/CP190 co-occupancy at topological boundaries (Figure 3A and B). Detailed examination of ∼800 annotated CG31365 peaks showed that CG31365 preferentially associates with transcription start sites (TSSs) or intragenic regions through recognition of a novel sequence motif, “ACTGCTT” (Figure S2C and 3C). Motifs for other CP190-interacting insulator proteins, Pita, CTCF, and Su(Hw), were also recovered (Figure S2D), suggesting that CG31365 can be recruited to the genome in combination with other insulator proteins. Consistent with this idea, k-means clustering revealed that CG31365 and CP190 peaks partially overlap with CTCF and Su(Hw) (Figure 3D and S2E). Overall, these results support the idea that CG31365 associates with a subset of topological boundaries in cooperation with CP190 and other insulator proteins.

**Figure 3.**
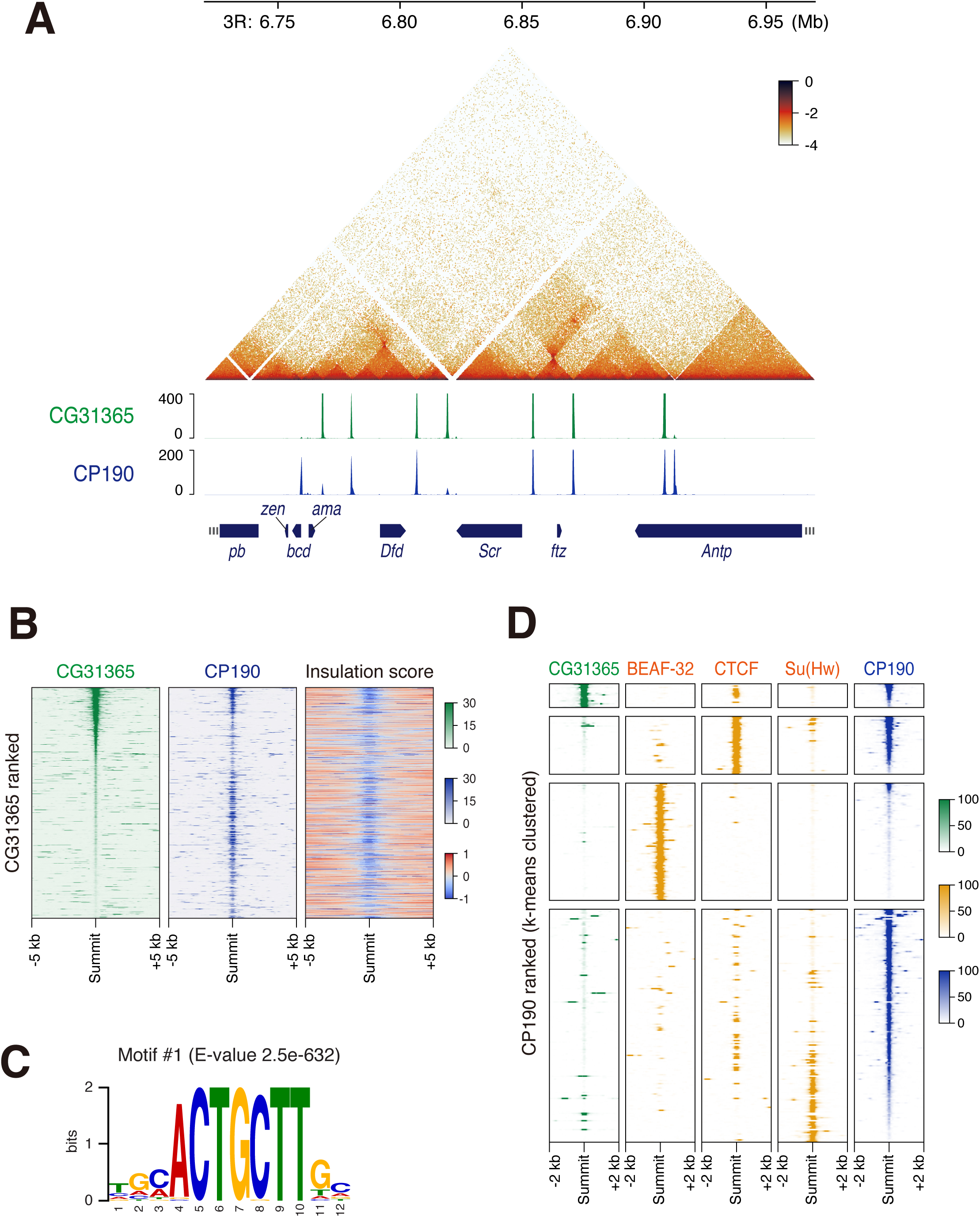
CG31365 occupies a subset of topological boundaries. (A) Micro-C contact-frequency map is shown alongside ChIP-seq tracks. Micro-C experiment was performed using *yw* embryos. Coolbox toolkit^79^ was used for visualization. (B) Heatmap visualization of ChIP-seq profiles at CG31365 peaks alongside insulation score calculated from *yw* Micro-C. Peaks are sorted by the CG31365 signal intensity. (C) A novel sequence motif at CG31365 peaks, annotated by MEME.^80^ This motif exhibited the lowest e-value relative to the others (see also Figure S2D). (D) Heatmap visualization of ChIP-seq profiles at CP190 peaks. Peaks are clustered by k-means (k = 5). Cluster 5 was omitted from visualization because it showed considerably small enrichment across all ChIP-seq profiles (see also Figure S2E). Peaks are sorted by the CP190 signal intensity within each cluster. For BEAF-32, CTCF, and Su(Hw), publicly available ChIP-seq tracks in nc14 embryos^36^ were used for the visualization. See also Figure S2.

### Chromatin association of CG31365 does not require ZAD-dependent condensation

To test whether the ZAD-dependent condensation activity of CG31365 is required for its chromatin association, GFP-3×FLAG-tagged WT or ΔZAD transgenes were individually introduced into the *CG31365[Δ1]* background (Figure 1C and 2E), and ChIP-seq was performed. Our analysis revealed that the genome-wide binding profile of CG31365 remains essentially unchanged even in the absence of the N-terminal ZAD (Figure 4A-C), indicating that CG31365 does not require ZAD-mediated condensation activity for recruitment to its chromatin-binding sites.

**Figure 4.**
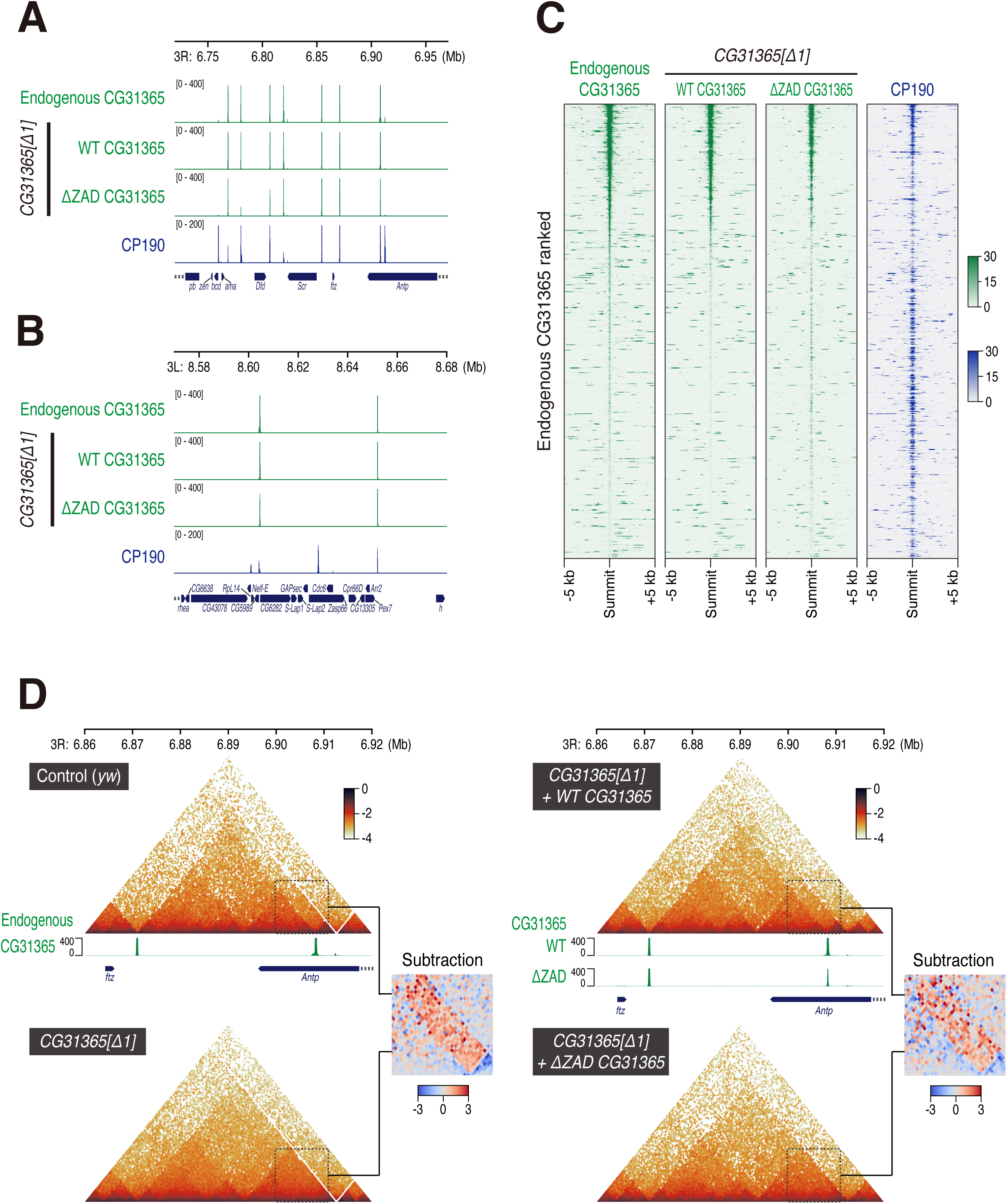
CG31365 regulates boundary formation through ZAD domain in a locus-specific manner. (A, B) Genome browser tracks visualized by IGV.^81^ For WT and ΔZAD CG31365, ChIP-seq experiments were performed using homozygous rescue strains (*CG31365[Δ1]* background). (C) Heatmap visualization of ChIP-seq profiles at endogenous CG31365 peaks. For WT and ΔZAD CG31365, ChIP-seq experiments were performed using homozygous rescue strains (*CG31365[Δ1]* background). (D) Micro-C contact-frequency maps are shown alongside ChIP-seq tracks. Micro-C experiments were performed using corresponding homozygous strains. ChIP-seq experiments were also performed using corresponding homozygous strains, except that *CG31365-GFP-3×FLAG* strain was used for endogenous CG31365. Differential contact profiles are summarized in subtraction heatmaps (*Mulb[Δ1]* − *yw* and *ΔZAD CG31365* rescue − *WT CG31365* rescue). Coolbox toolkit^79^ was used for visualization. See also Figure S3.

### CG31365 is required for boundary formation in a locus-specific manner

We next asked whether CG31365 contributes to boundary formation in developing embryos. To address this question, we performed Micro-C assays using embryos obtained from homozygous *CG31365[Δ1]* adults. This analysis revealed a loss of insulation activity specifically at a subset of CG31365-associated topological boundaries in mutant embryos (Figure 4D, left, and Figure S3A and S3B, left), suggesting that CG31365 plays an active role in regulating genome topology in a locus-specific manner. To further elucidate the role of ZAD-dependent condensation in this process, we performed additional Micro-C assays using embryos expressing either WT or ΔZAD CG31365 transgenes in the *CG31365[Δ1]* background. Intriguingly, the loss of insulation was restored by introducing the WT transgene (Figure 4D and S3, *CG31365[Δ1] + WT CG31365*), whereas the condensation-deficient ΔZAD mutant failed to rescue insulation despite being properly recruited to these loci (Figure 4D and S3, *CG31365[Δ1] + ΔZAD CG31365*). These findings indicate that CG31365 helps establish boundary formation in a locus-specific manner and that this function strictly depends on its ZAD-mediated condensation activity.

### CG31365/Mulberry associates with promoter-promoter loop anchors

It was recently proposed that 3D genome organization is regulated by two distinct classes of regulatory elements: topological boundaries and tethering elements.^3^ Intriguingly, our ChIP-seq analysis revealed that CG31365 is enriched not only at topological boundaries (Figure 3A and 3B) but also at the anchor regions of a long-range (∼80 kb) focal interaction connecting the promoters of two distant paralogous genes, *pyramus* (*pyr*) and *thisbe* (*ths*) (Figure 5A). *pyr* and *ths* encode FGF ligands closely related to the vertebrate FGF8/17/18 subfamily, and both genes display essentially identical expression patterns in the neurogenic ectoderm during mesoderm specification.^61^ At these anchor regions, CP190 was not enriched at CG31365 peaks (Figure 5A), suggesting that the association between CG31365 and CP190 is dependent on surrounding genomic context. We hereafter refer to CG31365 as Mulberry (Mulb), after the symbolic tree linking the tragic lovers Pyramus and Thisbe in Ovid’s *Metamorphoses*. The role of CG31365/Mulb in controlling long-range gene activation is discussed in the following sections.

**Figure 5.**
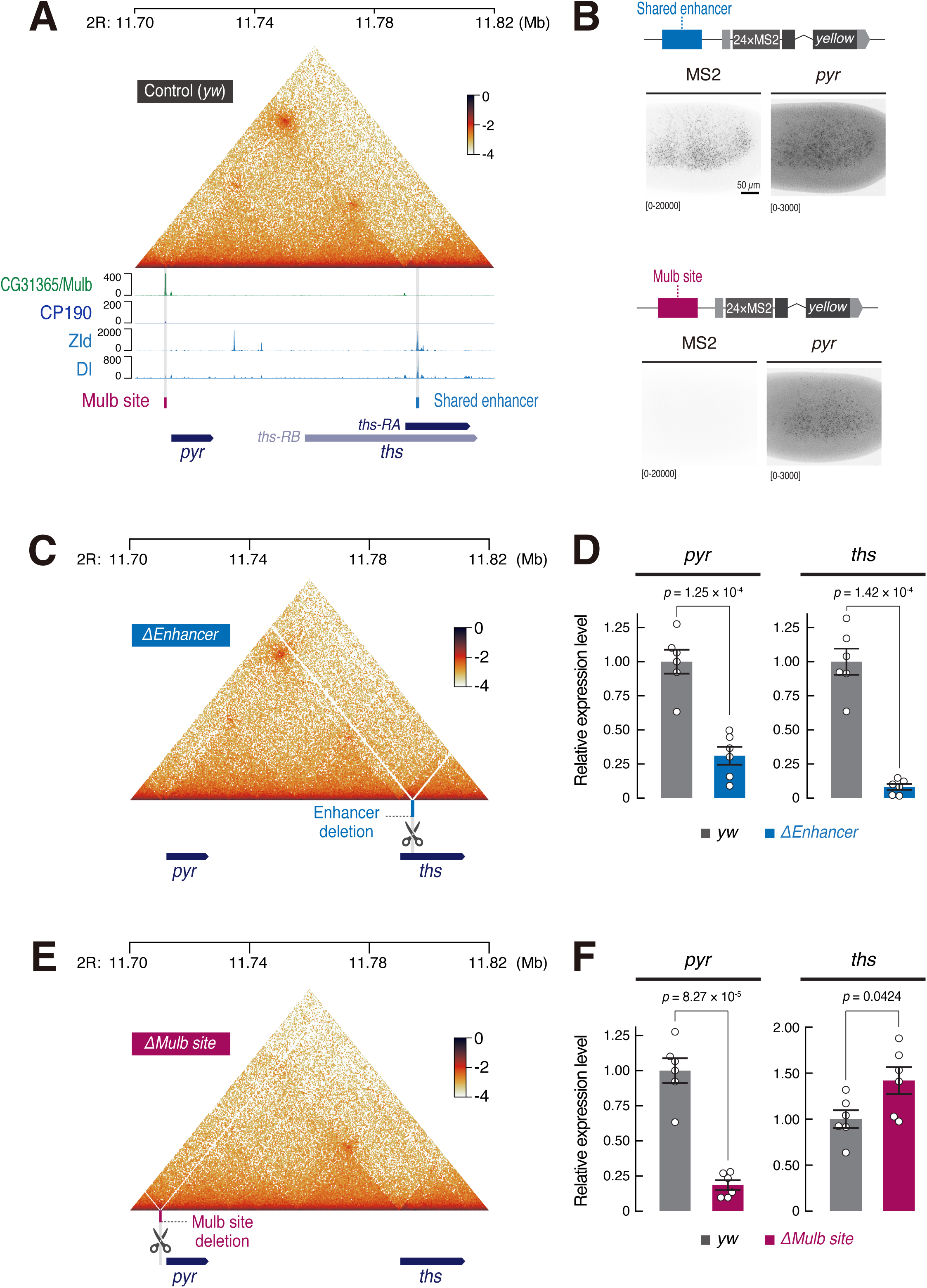
CG31365/Mulb associates with *pyr/ths* promoter-promoter loop. (A) Micro-C contact-frequency map is shown alongside ChIP-seq tracks. Micro-C experiment was performed using *yw* embryos. Publicly available ChIP-seq tracks for Zld in 120-150 min AEL embryos (GEO: GSE30757)^82^ and Dl in 2-4 h AEL embryos (GEO: GSE55306)^83^ were used for visualization. In the gene track, dark- and dim-colored isoforms represent active and inactive isoforms in nc14 embryos, respectively (see also Figure S4A for Pol II ChIP-seq track). Coolbox toolkit^79^ was used for visualization. (B) Schematic representations of *24×MS2-yellow* transgenes, alongside representative smiFISH images of embryos carrying the corresponding transgenes. Maximum projected images were rotated to display the anterior side left and posterior right. (C) Micro-C contact-frequency map in homozygous *ΔEnhancer* embryos, lacking the shared enhancer at *pyr*/*ths* locus. Coolbox toolkit^79^ was used for visualization. (D) Relative expression levels of *pyr* and *ths* measured by qRT-PCR. Expression levels were normalized to *RpL32* and scaled to *yw* embryos. Bars represent the mean, and error bars represent the standard error of the mean. The *p*-values were calculated using a two-sided Welch’s *t*-test. n = 6 for *yw* embryos; n = 6 for homozygous *ΔEnhancer* embryos. (E) Same as (C), but homozygous *ΔMulb site* embryos, lacking the Mulb enrichment site at the promoter region of *pyr*, were used. (F) Same as (D), but *yw* and homozygous *ΔMulb site* embryos were used. n = 6 for *yw* embryos; n = 6 for homozygous *ΔMulb site* embryos. For *yw*, the data is the same as that shown in Figure 5D. See also Figure S4.

### Mulb-anchored loop fosters co-regulation of two distant genes by a shared enhancer

In early embryos, transcription of the *ths* short isoform (*ths-RA*) and distal *pyr* is thought to be co-regulated by a shared enhancer located within the intronic region of the *ths* gene (Figure 5A and S4A).^61,62^ When tested in isolation, this putative shared enhancer was able to drive reporter gene transcription specifically in regions where endogenous *pyr* is active (Figure 5B). In contrast, no reporter expression was detected when the Mulb site upstream of the *pyr* promoter was used (Figure 5B), indicating that Mulb does not function as a direct transcriptional activator.

Having confirmed this, we next examined the biological function of these two DNA elements in the endogenous *pyr*/*ths* locus. We first removed the putative shared enhancer from the *ths* intron using CRISPR/Cas9-mediated genome-editing. Micro-C analysis of the resulting mutant embryos showed that the long-range (∼80 kb) promoter-promoter loop remained essentially intact despite removal of the intronic enhancer (Figure 5C). This result indicates that formation of the long-range focal loop does not depend on the presence of a nearby active enhancer. Importantly, although the promoter-promoter loop was preserved, enhancer deletion caused a significant reduction in expression of both the enhancer-proximal *ths* and the distal *pyr* gene (Figure 5D), providing direct evidence that the enhancer is shared by two distant promoters. We then removed the Mulb site upstream of the *pyr* promoter using the same genome-editing strategy. In contrast to the enhancer-deletion assay (Figure 5C), Micro-C analysis revealed a clear loss of the long-range promoter-promoter loop (Figure 5E). Intriguingly, in this rearranged genomic configuration, expression of the distal *pyr* gene was selectively abolished (Figure 5F). Together, these results support a model in which a Mulb-anchored promoter-promoter loop enables co-regulation of two distant genes by a shared enhancer. Notably, we reproducibly observed upregulation of the enhancer-proximal *ths* gene in the absence of promoter-promoter looping (Figure 5F), suggesting that a promoter-competition mechanism operates to balance expression levels of functionally related genes.^63,64^ Supporting this view, deletion of the *ths* promoter (*Δths TSS*) eliminated *ths* expression while partially increasing distal *pyr* expression (Figure S4). These findings are consistent with the presence of cross-regulatory mechanisms between functionally related distant genes at other developmental loci in *Drosophila* (see Discussion).^31^

### Mulb condensates foster the formation of long-range focal contacts

We next sought to elucidate the molecular mechanism underlying Mulb-mediated long-range loop formation. To this end, we used single-molecule RNA FISH and immunofluorescence to simultaneously visualize Mulb condensates and the active transcription sites of *pyr* and *ths* in nc14 embryos. Airyscan super-resolution imaging revealed frequent colocalization between Mulb condensates and *pyr*/*ths* transcription sites (Figure 6A and 6B). This observation was further supported by average profiling of active *pyr* and *ths* transcription sites (Figure 6C and 6D). Notably, such clear colocalization was not observed at the Mulb-depleted *even-skipped* (*eve*) locus (Figure 6C, 6D, and S5), indicating that Mulb condensates preferentially associate with their target genomic regions.

**Figure 6.**
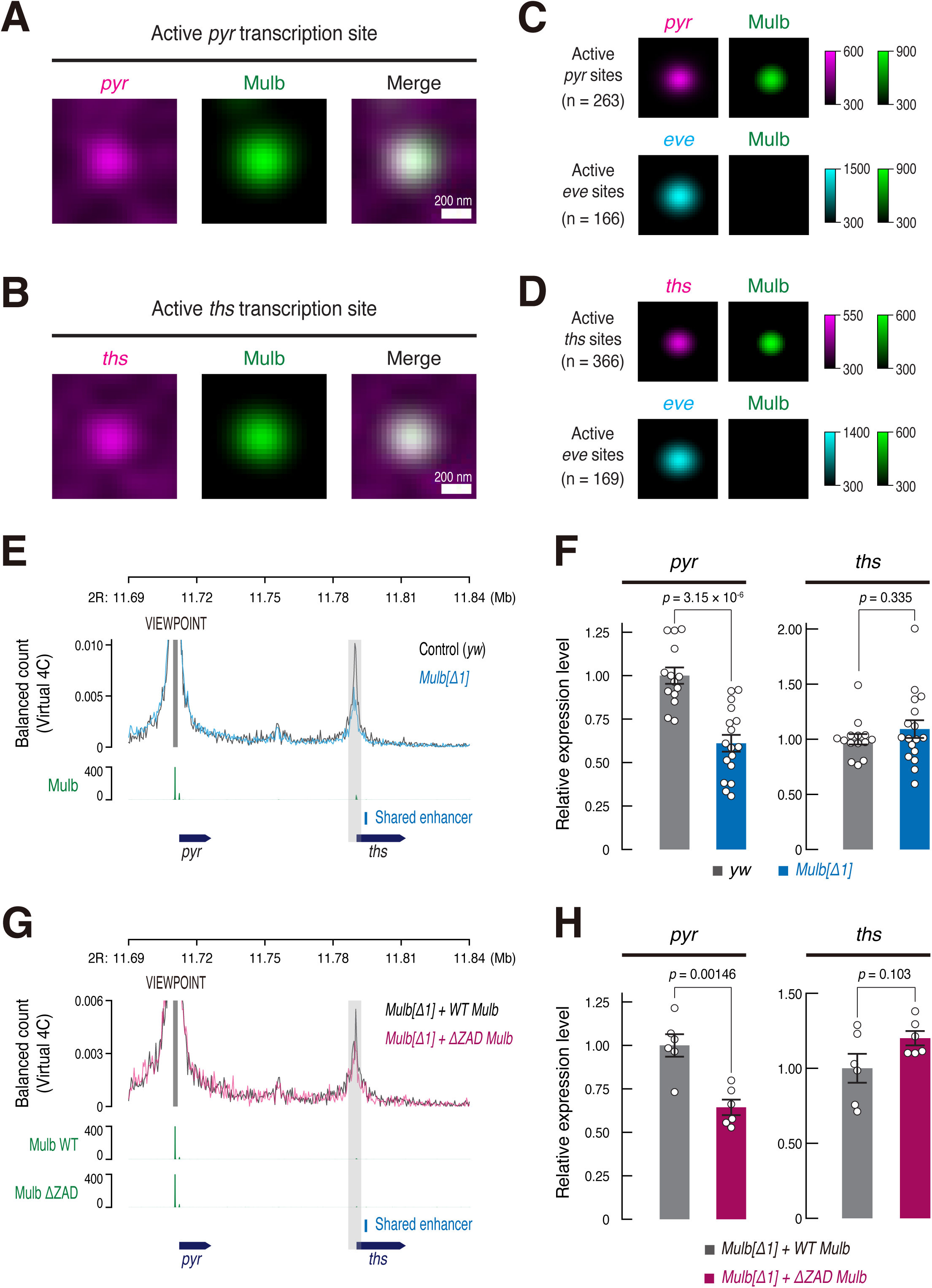
Mulb condensates foster the formation of *pyr/ths* promoter-promoter loop. (A) Representative *pyr* and Mulb signals at an active *pyr* transcription site. Active transcription sites were determined based on foci brightness and nuclear localization (see Method Details). A single Airyscan z-slice image was cropped around an active *pyr* transcription site. *pyr* smiFISH signal and Mulb staining signal are shown in magenta and green, respectively. (B) Same as (A), but an active *ths* transcription site was analyzed. (C) Averaged profiles of Mulb distribution centered around (top) *pyr* and (bottom) *eve* transcription sites. Five independent embryos were analyzed. The counts of analyzed transcription sites are indicated in the figure. (D) Same as (C), but averaged profiles of Mulb distribution centered around (top) *ths* and (bottom) *eve* are shown. Six independent embryos were analyzed. (E) Virtual 4C profiles in *yw* and homozygous *Mulb[Δ1]* embryos alongside ChIP-seq track of endogenous Mulb. For the viewpoint, a 2 kbp-scaled anchor of *pyr*/*ths* promoter-promoter loop^3^ (2R:11709600-11711600) was used. Region around *ths* promoter is highlighted. (F) Relative expression levels of *pyr* and *ths* measured by qRT-PCR. Expression levels were normalized to *RpL32* and scaled to *yw* embryos. Bars represent the mean, and error bars represent the standard error of the mean. The *p*-values were calculated using a two-sided Welch’s *t*-test. n = 14 for *yw* embryos; n = 17 for homozygous *Mulb[Δ1]* embryos. (G) Same as (E), but Virtual 4C profiles in homozygous rescue embryos (*Mulb[Δ1]* background) are shown alongside ChIP-seq tracks of WT and ΔZAD Mulb. (H) Same as (F), but homozygous rescue embryos (*Mulb[Δ1]* background) were used. Expression levels were scaled to homozygous *WT Mulb* rescue embryos. n = 6 for homozygous *WT Mulb* rescue embryos; n = 6 for homozygous *ΔZAD Mulb* rescue embryos. See also Figure S5, S6, and S7.

To further investigate the functional significance of Mulb association at the *pyr/ths* locus, we performed virtual 4C analysis using Micro-C datasets from control and *Mulb[Δ1]* embryos. Loss of Mulb resulted in a marked reduction in promoter-promoter loop strength (Figure 6E). Consistent with the Mulb-site deletion experiment (Figure 5E), weakening of the promoter-promoter loop selectively reduced expression of the distal *pyr* gene (Figure 6F), supporting the idea that Mulb organizes 3D genome topology to facilitate “enhancer sharing” between two distant genes. We then conducted the same analyses using Micro-C data from embryos expressing WT or ΔZAD Mulb in the *Mulb[Δ1]* background. Importantly, both the reduction of promoter-promoter loop strength (Figure 6G) and the selective loss of distal *pyr* expression (Figure 6H) were similarly observed upon loss of the ZAD-dependent condensation activity of Mulb (Figure 2E). Misregulation of another Mulb-associated pair of distant (∼70 kb) paralogous genes, *disco* and *disco-related* (*disco-r*), was also observed in these mutants (Figure S6). We therefore concluded that Mulb condensates play an active role in facilitating the coordinated expression of functionally related distant genes by organizing 3D genome topology in a locus-specific manner (Figure 7). This mechanism also appears to operate at other non-paralogous loci (Figure S7), underscoring the generality of our findings.

**Figure 7.**
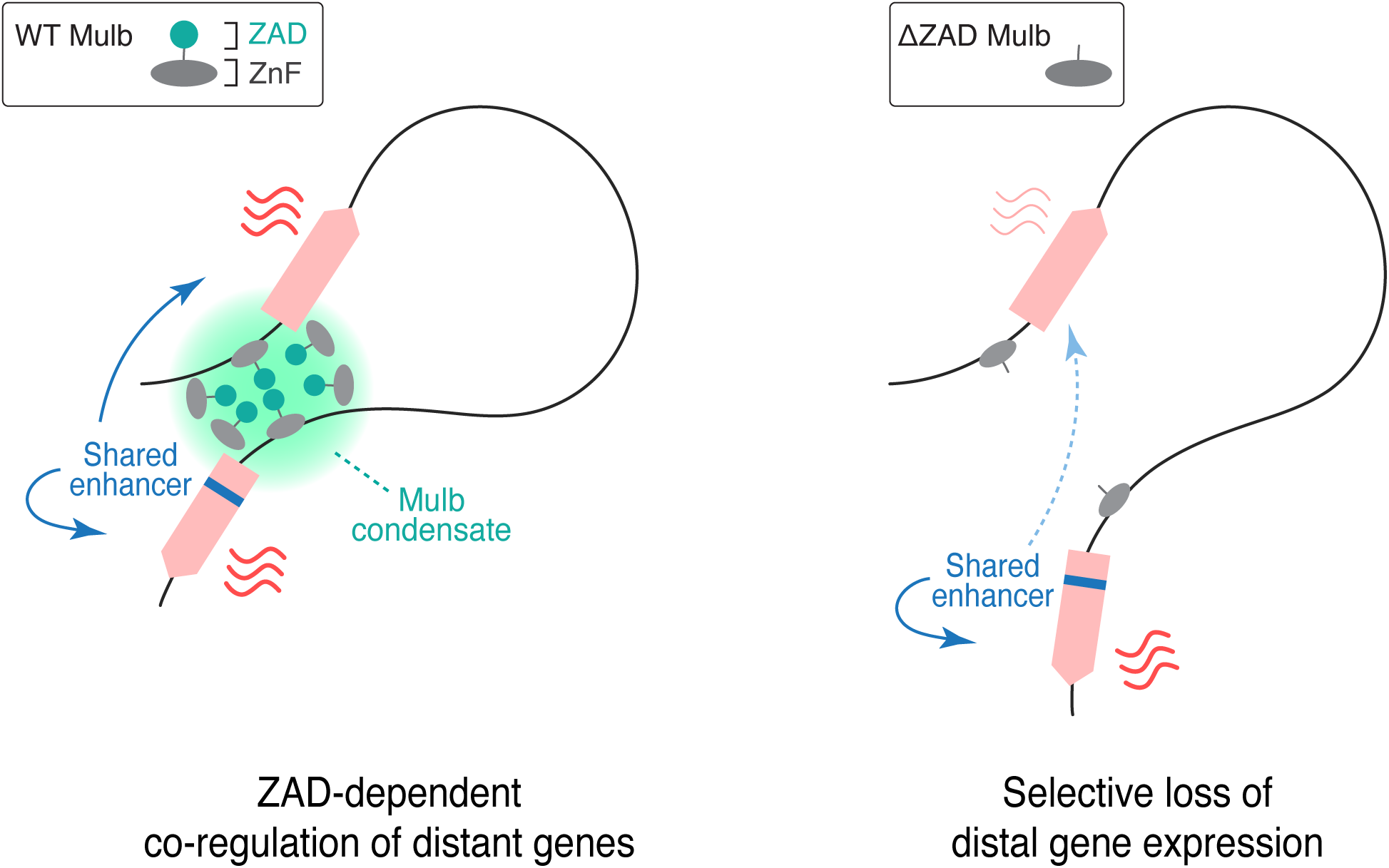
Role of Mulb in coordinating the expression of functionally related distant genes. Mulb condensates facilitate the co-regulation of distant genes, whereas the expression of the distal gene is selectively lost upon the loss of condensation activity in the ΔZAD mutant embryos.

## Discussion

### Mulb as a condensate-dependent regulator of 3D genome organization

In this study, we identified the ZAD-ZnF protein Mulb as a new regulator of 3D genome organization in *Drosophila*, facilitating both topological insulation and long-range looping interaction in developing embryos. A key molecular feature of Mulb is its ZAD-dependent condensation activity (Figure 2E). Similar to Mulb, the looping factor GAF is also known to form nuclear condensates.^58,65–67^ However, GAF condensation relies on its binding to AAGAG satellite repeats through its C-terminal DNA-binding domain, rather than on self-association mediated by the N-terminal POZ/BTB domain.^67^ Functionally, condensates formed by GAF and Mulb also display distinct molecular properties. A recent study reported that GAF condensates co-localize with HP1 to silence transcription from AAGAG satellite repeats in early embryos.^67^ In contrast, Mulb exerts its genome-organizing function through the assembly of ZAD-dependent condensates that facilitate long-range gene activation by physically associating with its target loci (Figure 2, 4, 6). Importantly, previous studies suggest that similar condensation-dependent mechanisms also operate in other biological contexts. For example, dynamic rearrangements of neighboring TADs are mediated by Oct4 condensation during the reprogramming of mouse embryonic fibroblasts.^68^ Similarly, during the yeast heat-shock response, Hsf1 condensates have been proposed to foster coalescence of Hsf1-targeted heat-shock-induced genes.^69^ In addition, recent biophysical studies suggest that DNA-protein co-condensation can generate mechanical forces that can bring distant genomic loci into close proximity.^70–73^ Taking these findings into consideration, one possible explanation is that Mulb-mediated genome organization involves the generation of mechanical forces that pull distant genomic loci together via condensate formation. In mammals, CTCF has also been shown to self-associate and form clusters during the formation of TADs.^74^ At this point, we cannot rule out the possibility that ZAD-mediated self-association is the primary mechanism driving Mulb-dependent loop formation, with Mulb condensates forming as a secondary. Clearly, future studies will be required to further elucidate the molecular function of Mulb, including the identification of its potential interacting partners.

### Mulb organizes multi-way regulatory hub in developing embryos

By combining Micro-C/ChIP-seq assays and targeted genome-editing, we demonstrated that Mulb facilitates the formation of multi-way regulatory hub at the *pyr*/*ths* locus, thereby enabling “enhancer sharing” by two distant (∼80 kb) promoters (Figure 7). This conclusion is nicely illustrated by our CRISPR deletion of the Mulb binding site upstream of the *pyr* promoter, which resulted in both the loss of long-range looping interactions (Figure 5E) and the selective reduction of distal *pyr* expression (Figure 5F). Under this genomic configuration, we reproducibly observed upregulation of the enhancer-proximal *ths* gene (Figure 5F), suggesting a cross-regulatory mechanism that balances the expression of functionally related genes over large genomic distances.^31^ Intriguingly, full-locus deletion of *Mulb* did not result in a complete loss of long-range looping interactions at the *pyr*/*ths* locus (Figure 6E), and the reduction in distal *pyr* expression was correspondingly milder (Figure 6F) compared with the CRISPR deletion of the Mulb binding site (Figure 5E and F). These observations suggest that the upstream anchor region of *pyr* likely associates with additional, yet unidentified factor(s) to mediate looping interaction. Given our recent finding that ZAD-ZnFs broadly exhibit intrinsic insulator-binding activity in early embryos,^57^ we speculate that Mulb-dependent loops and topological insulation are combinatorially regulated with other proteins. While the present study focused on early embryogenesis, the presence of ultra-long-range “meta-loops” in the *Drosophila* brain^29^ raises the possibility that Mulb may also contribute to long-range looping interactions in other tissues such as neurons. These possibilities warrant future investigation.

### ZAD-dependent genome-organization activity of Mulb

*Mulb* is a member of the evolutionarily dynamic ZAD-ZnF gene family, which has undergone rapid expansion during insect evolution.^40^ Among this gene family, we found that *Mulb* is conserved across divergent *Drosophila* species, not only at the level of amino acid composition but also in the dimerization activity of its N-terminal ZAD (Figure S1), suggesting that Mulb-mediated loop formation is an ancient mechanism that likely existed in their common ancestor. Importantly, previously identified insulator proteins such as M1BP, Pita, ZIPIC, and Zw5 are also members of the ZAD-ZnF protein family in *Drosophila*.^48,53,75^ More recently, our systematic whole-genome analysis demonstrated that many ZAD-ZnFs other than these factors also exhibit intrinsic insulator-binding activity.^57^ Critically, however, the molecular function of the N-terminal ZAD has not been examined so far in the context of *Drosophila* development. Our finding that Mulb exerts its genome-organizing function through the ZAD domain provides key mechanistic insight into the fundamental principles of 3D genome organization as well as evolutionary origins of ZAD-ZnFs in *Drosophila*. Together with POZ/BTB-containing architectural proteins such as GAF, Lola-I, CP190, and Mod(mdg4),^13,28–30,76^ we propose that ZAD-ZnFs constitute a previously unrecognized class of genome organizers in *Drosophila* and other insect species.

## Acknowledgement

We thank Hitomi Takishita and Misako Sato for fly husbandry, and the Bloomington *Drosophila* Stock Center for fly strains. We are also grateful to Ryuichiro Nakato for support on the RNA-seq analysis, Yuka W. Iwasaki and Yusuke Kishi for advice on the ChIP-seq experiment, Chikara Takeuchi for sharing a Micro-C protocol, and members of the Fukaya laboratory for critical comments on the manuscript. T.F. was supported by FOREST (JPMJFR214W) and CREST program (JPMJCR25T2) from the JST (Japan Science and Technology Agency), a Grant-in-Aid for Scientific Research (A) (25H00967), a Grant-in-Aid for Transformative Research Areas (A) (24H02327) from the JSPS (Japan Society for the Promotion of Science), and the research grant from the Takeda Science Foundation. Y.U. was supported by JSPS fellowship (24KJ0843).

## Author contributions

Y.U. and T.F. designed and conceived this study. Y.U. performed all the experiments and data analysis. T.F. supervised the project. Y.U. and T.F. wrote the manuscript. All the authors approved the manuscript.

## Key resources table

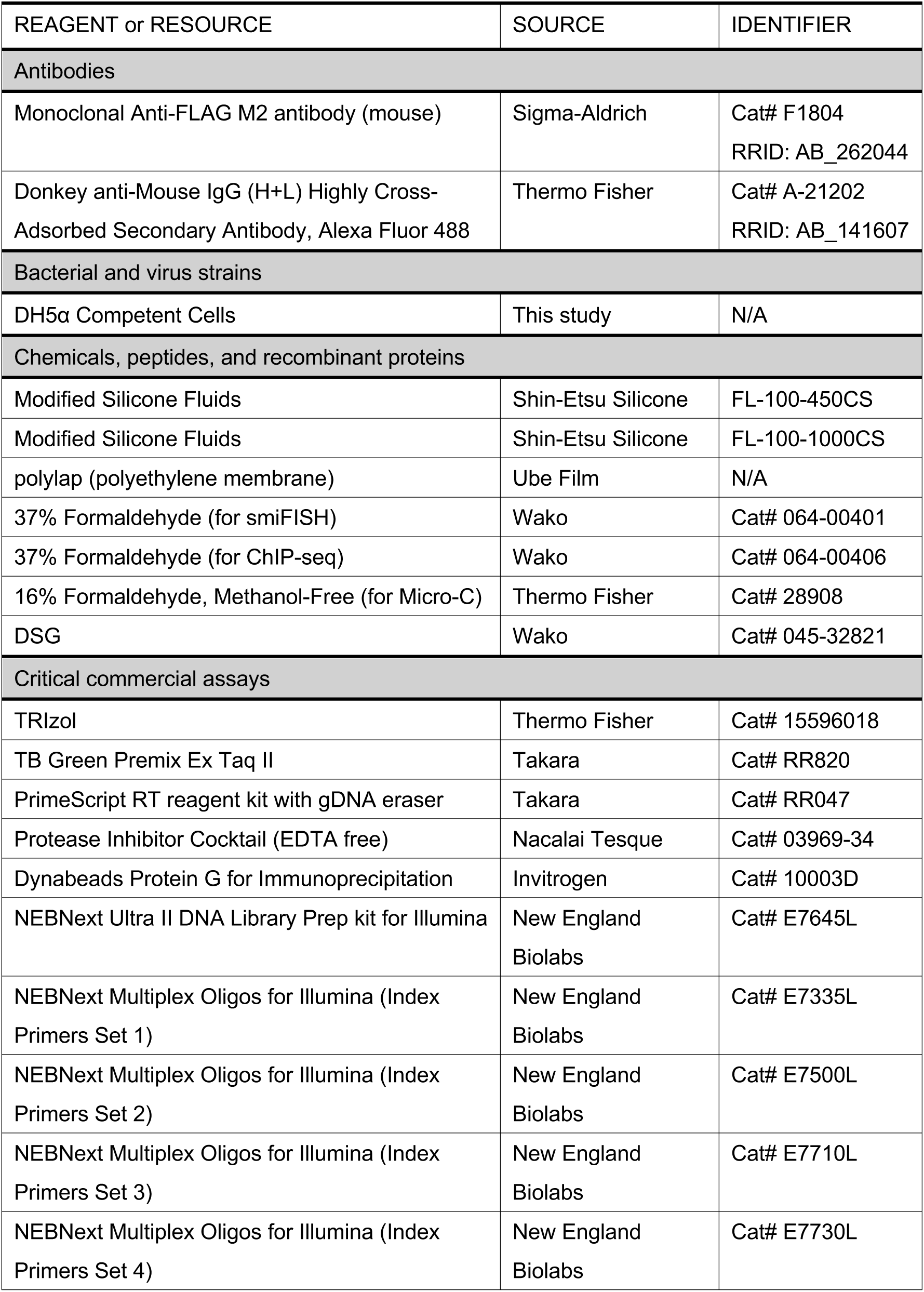
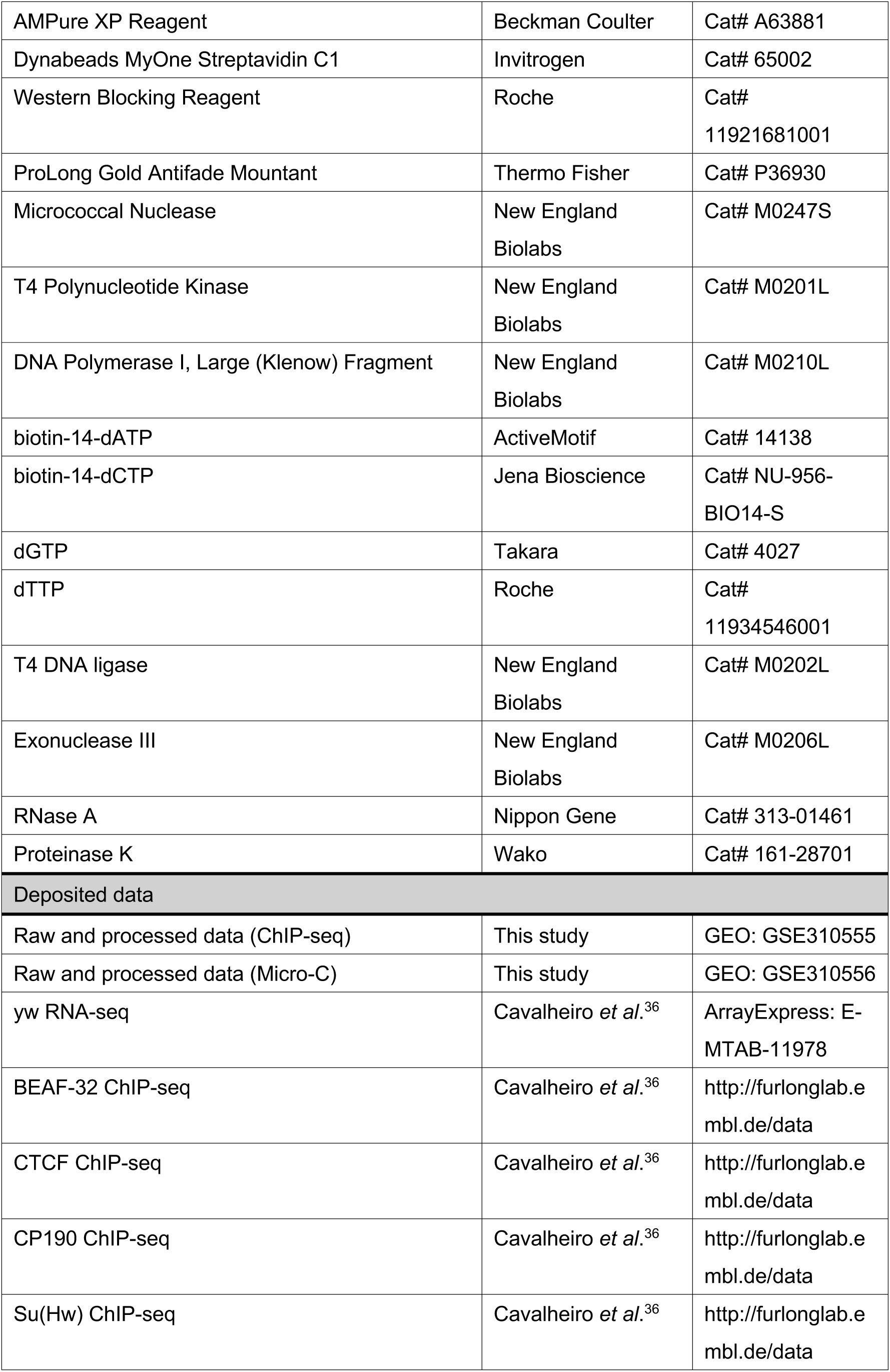
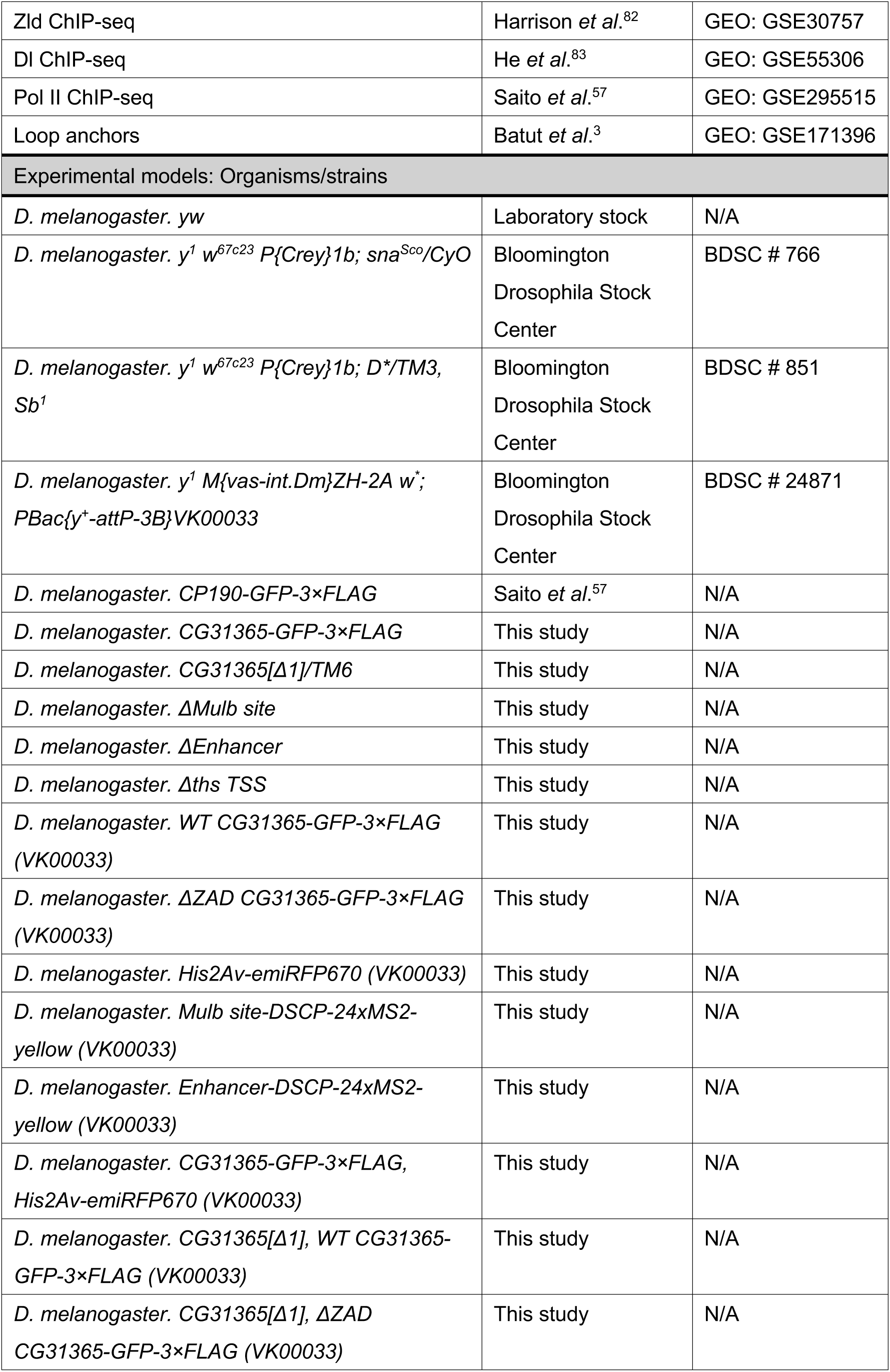
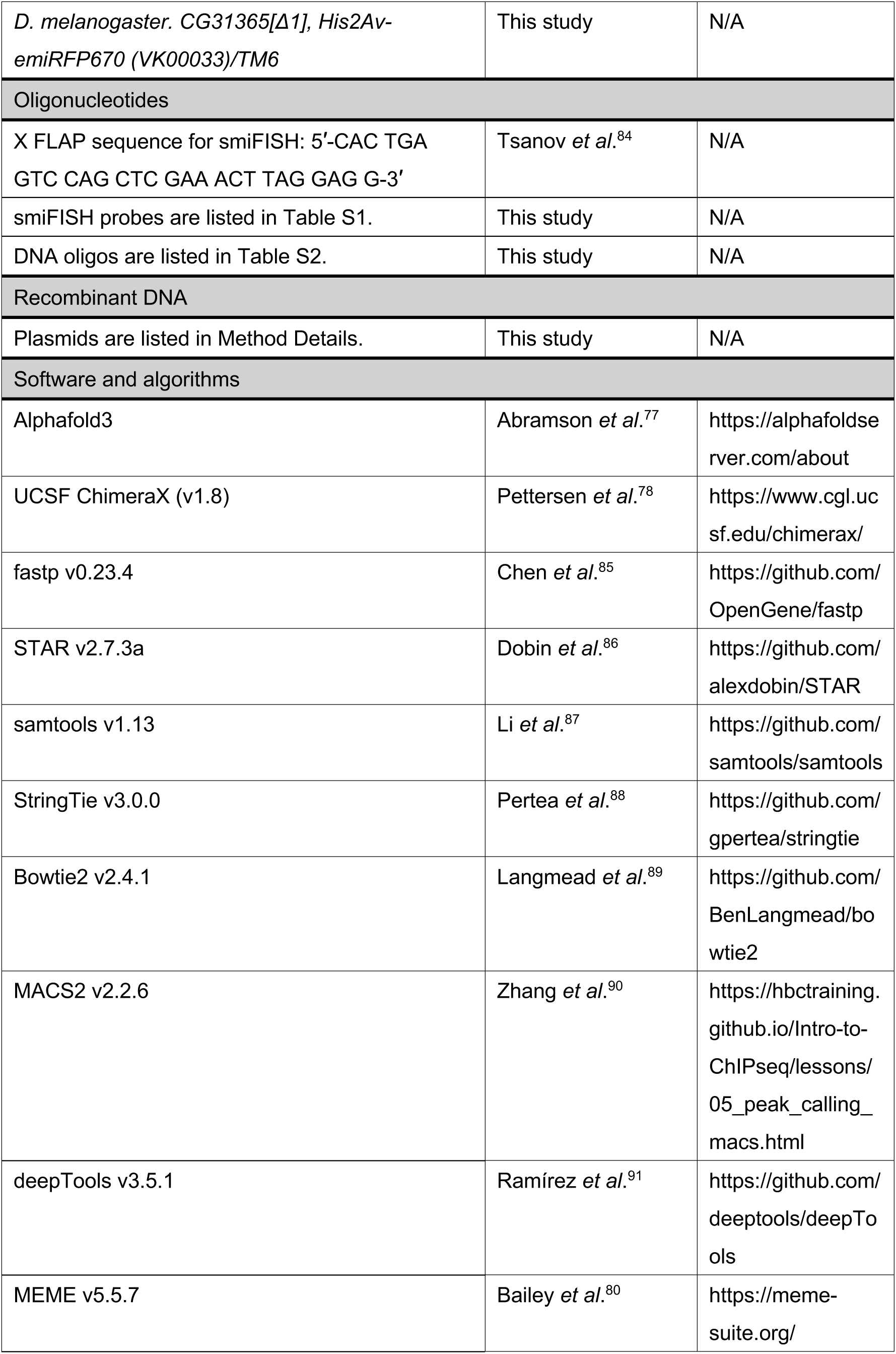
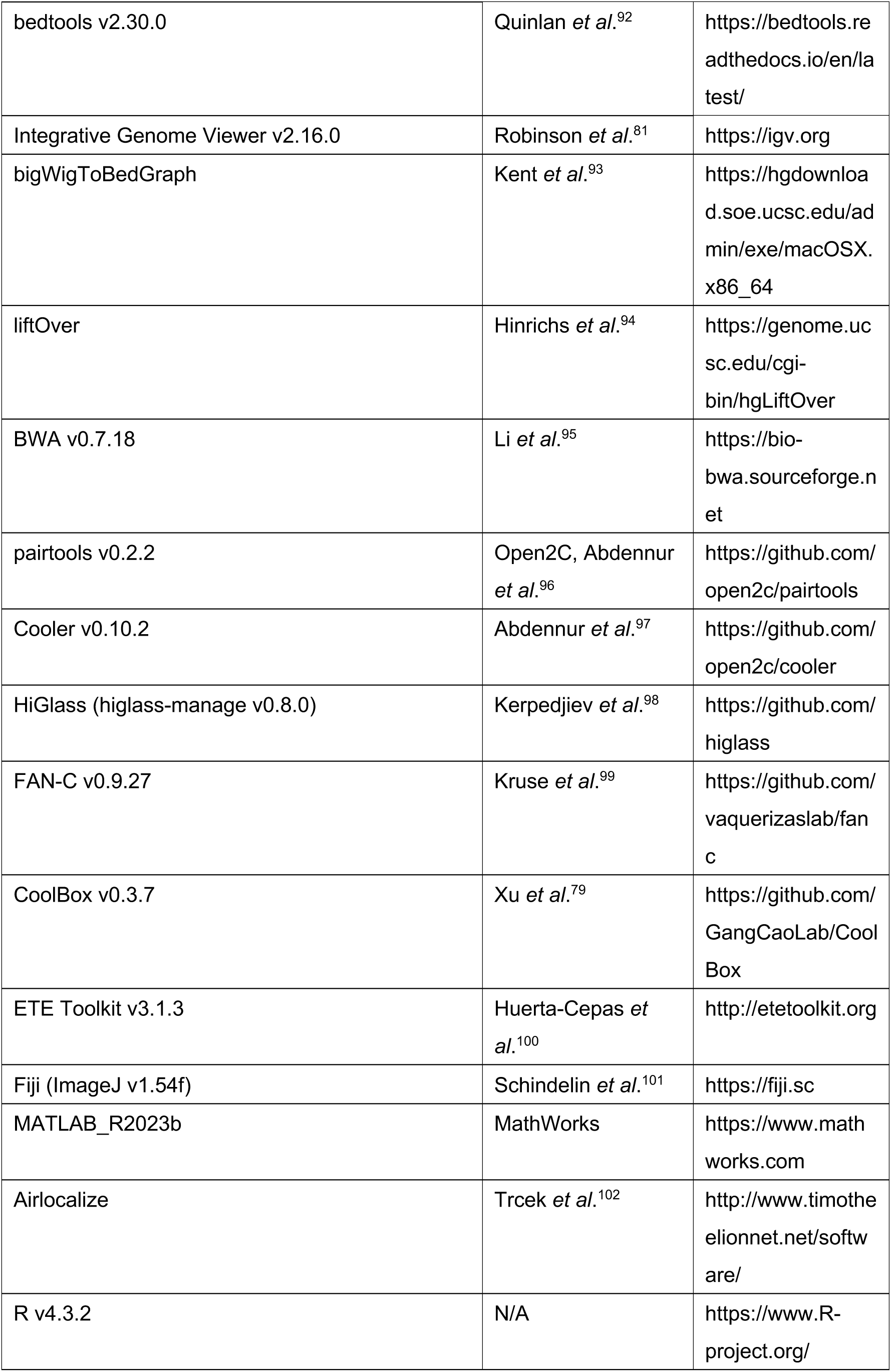
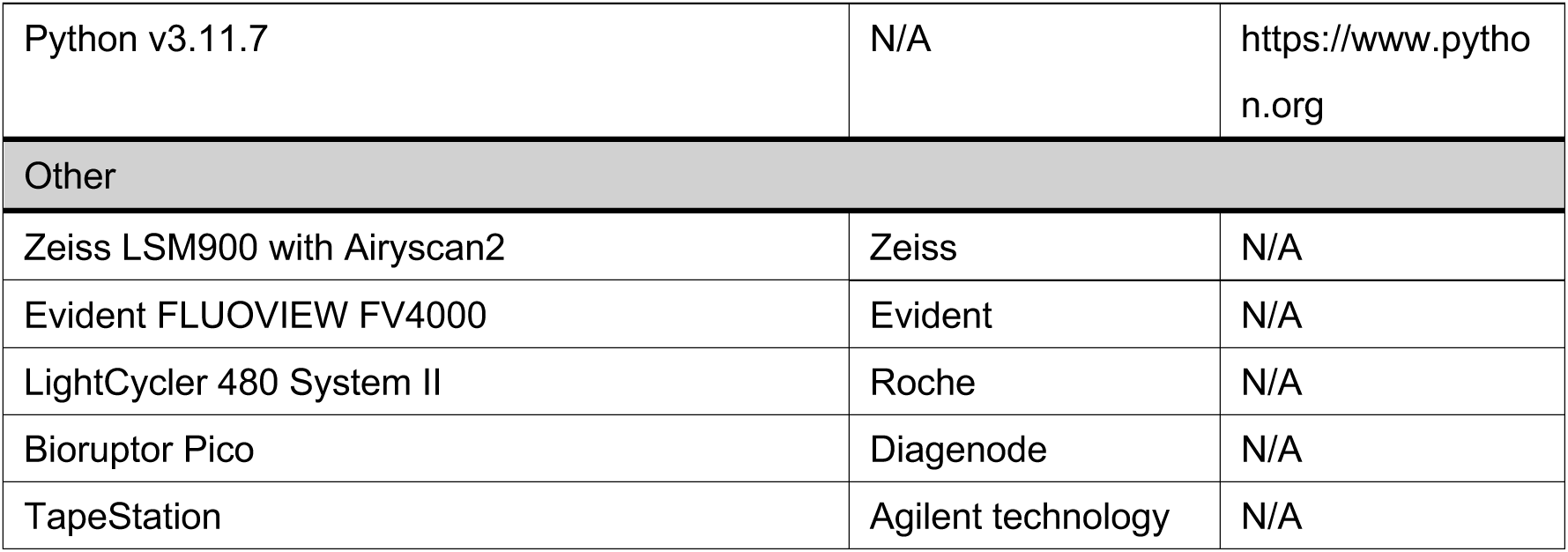

## Data Availability

All sequencing data have been deposited in the Gene Expression Omnibus under accession code GSE310555 (ChIP-seq) and GSE310556 (Micro-C).

## Experimental model and study participant details

All *Drosophila melanogaster* lines used in this study are listed in Key resources table. Detailed constructions of each fly line are described below.

## Method Details

### CRISPR/Cas9-mediated genome-editing of endogenous loci

Several genome-edited fly lines were obtained using CRISPR/Cas9-mediated homology directed repair. To generate *CG31365[Δ1]* strain, two gRNAs were designed to target PAM sequences flanking the open reading frame of *CG31365*. A pair of pCFD3 gRNA expression plasmids, pBS-hsp70-Cas9 plasmid (addgene #46294) and pBS-loxP-3×P3-dsRed-loxP donor plasmid containing 5′ and 3′ homology arms were co-injected using *yw* strain. Injection mixture contained 0.2 µg/µL of each gRNA expression plasmid, 0.8 µg/µL of donor plasmid, 0.6 µg/µL of Cas9 expression plasmid, 5 mM KCl and 0.1 mM phosphate buffer (pH 6.8). Microinjection was performed as previously described.^103^ In brief, 0-1 h embryos were collected and dechorionated with bleach. Aligned embryos were dried with silica gel for ∼8 min and covered with FL-100-1000CS silicone oil (Shin-Etsu Silicone). Subsequently, microinjection was performed using FemtoJet (Eppendorf) and DM IL LED inverted microscope (Leica) equipped with M-152 Micromanipulator (Narishige). *3×P3-dsRed* marker was used for screening.

To generate *CG31365-GFP-3×FLAG* strain, a gRNA was designed to target PAM sequence just upstream of the stop codon. pCFD3 gRNA expression plasmid, pBS-hsp70-Cas9 plasmid (addgene #46294) and pBS-GFP-3×FLAG-loxP-3×P3-dsRed-loxP donor plasmid containing 5′ and 3′ homology arms were co-injected using *yw* strain. Microinjection was performed as described above. *3×P3-dsRed* marker was used for screening. Resulting genome-edited flies were then crossed with *y^1^ w^67c^*^23^ *P{Crey}1b; D^∗^/TM3, Sb^1^* (BDSC# 851) to remove *3×P3-dsRed* marker from the endogenous locus.

To generate *ΔMulb site*, *ΔEnhancer* and *Δths TSS* strains, two gRNAs were designed for each strain to target PAM sequences flanking the regions to be deleted. A pair of pCFD3 gRNA expression plasmids, pBS-hsp70-Cas9 plasmid (addgene #46294) and pBS-loxP-3×P3-GFP-loxP donor plasmid containing 5′ and 3′ homology arms were co-injected using *yw* strain. Injection mixture contained ∼0.4 µg/µL of each gRNA expression plasmid, ∼0.8 µg/µL of donor plasmid, ∼0.6 µg/µL of Cas9 expression plasmid, 5 mM KCl and 0.1 mM phosphate buffer (pH 6.8). Microinjection was performed as described above. *3×P3-GFP* marker was used for screening. Resulting genome-edited flies were then crossed with *y^1^ w^67c^*^23^ *P{Crey}1b; sna^Sco^/CyO* (BDSC# 766) to remove *3×P3-GFP* marker from the endogenous locus.

### Site-specific transgenesis by phiC31 system

GFP-3×FLAG fused WT and ΔZAD CG31365 transgenes, His2Av-emiRFP670 transgene, Mulb site-DSCP-24×MS2-yellow transgene and Enhancer-DSCP-24×MS2-yellow transgene were individually integrated into a unique *attP* landing site on the third chromosome using VK00033 strain.^104^ Injection mixture contained ∼1.0 µg/µL of plasmid DNA, 5 mM KCl and 0.1 mM phosphate buffer (pH 6.8). Microinjection was performed as described above. For His2Av-emiRFP670 transgene, previously constructed pbφ-His2Av-emiRFP670 plasmid^57^ was used.

### Scoring of embryo hatching rates

More than 120 virgin females and males were individually collected for 2 days, mated with each other, and aged for 2 additional days. Following two ∼1-hour pre-lays, embryos were collected for 3 hours and incubated on apple juice plates for ∼48 hours at 25 °C. Subsequently, the number of unhatched embryos was counted to calculate the embryo hatching rate, expressed as the percentage of hatched embryos from total. Each replicate consisted of a batch of 100 embryos.

### RNA extraction from embryos

More than 70 virgin females and males were individually collected for 1-2 days, mated with each other, and aged for a day. For the collection of nc14 embryos, after two ∼1-hour pre-lays, embryos were collected over 30 min and aged for 2 h 10 min at 25 °C. Embryos were then immersed in FL-100-450 (Shin-Esu Silicone) and those corresponding to nc14 were hand-sorted using a stereo microscope (LEICA M205C). For qRT-PCR analysis of *disco* and *disco-r*, 8–10 h after egg laying (AEL) embryos were collected following the same procedures, except that embryos were collected over 2 h, aged for 8 h, and hand-sorted to collect embryos without severe developmental defects. Subsequently, embryos were immersed in TRIzol reagent (Thermo Fisher) and homogenized with pestles. The resulting lysates were snap-frozen in liquid nitrogen and stored at -80 °C until use, or immediately processed for RNA extraction. For each replicate, approximately 5-10 embryos were used. Total RNA was extracted using TRIzol reagent, following the manufacturer’s protocol.

### qRT-PCR

Reverse transcription was performed using PrimeScript RT reagent kit with gDNA eraser (Takara), following the manufacturer’s protocol. For each replicate, 100 ng of total RNA was used. qRT-PCR was performed using TB Green Premix Ex Taq II (Takara) and the LightCycler 480 System II (Roche). For each measurement, technical triplicates were prepared. Serial dilutions of cDNA were used for the calibration. Melting-curve analysis was performed to confirm the amplification of a single product for each target. For *pyr* and *ths*, the expression of *opa* gene was measured to confirm that replicates were enriched for the same developmental stage in nc14. Measurements for *disco* and *disco-r* were performed with 8-10 h AEL embryos because *disco*/*disco-r* is not coactivated in nc14 embryos.^105^ The expression of *RpL32* was measured as an internal control. The same set of RNA samples was used in each experiment for *pyr*/*ths* or *disco*/*disco-r* gene pairs. The following primer pairs were used for the analysis. *RpL32*: (5′-CCG CTT CAA GGG ACA GTA TCT G-3′) and (5′-ATC TCG CCG CAG TAA ACG C-3′). *pyr*: (5′-CTG CTA TTG CGT GGT CAG C-3′) and (5′-GTA TGT TCT CGG CGG TCA CT-3′). *ths*: (5′-GTG GTG ATT TCA GGT GCA TT-3′) and (5′-AGA CTG TTC CTT CCG CAG CTA-3′). *opa*: (5′-TCC AGC GTT AAG CAG GAG AT-3′) and (5′-CTC GTG CAT CGA GTG GAA TA-3′). *disco*: (5′-CTC CCA TCT ACC GTC ACA CC-3′) and (5′-AAG GAT GAA CCT GAT CCA GTG-3′). *disco-r*: (5′-GAA TAT TGG CCA CGA GAA GC-3′) and (5′-GGC GGC TAA ACG ATG AAC T-3′).

### Embryo collection and fixation for ChIP-seq and Micro-C

To collect homozygous *CG31365[Δ1]* mutant embryos, homozygous males and homozygous virgin females from *CG31365[Δ1]/TM6* strain were collected over a period of three days. Collected flies were transferred to population cages containing apple juice agar plates supplemented with yeast paste and allowed to mate for 1 day at 25 °C. Embryos were collected over the next two days. For all other strains, aged adult flies were discarded from vials in the morning, and newly eclosed flies were collected in the next evening. These flies were similarly transferred to population cages with apple juice agar plates supplemented with yeast paste and allowed to mate for 1 day at 25 °C, and embryos were collected on the following day. On the day of embryo collection, after two ∼1-hour pre-lays, embryos were collected over 30 min and aged for 2 h 10 min at 25 °C. This resulted in collection of 2h10-2h40 AEL embryos, corresponding to nc14. Collected embryos were dechorionated by immersion in bleach for 2 min, followed by thorough rinsing with water. Embryos were then subjected to the fixation process. For ChIP-seq, embryo fixation was performed as described in Saito *et al*.^57^ Embryos were fixed by rotating in a mixture of heptane and PBST [1× PBS and 0.5% TritonX-100] containing 1.8% formaldehyde (Wako; final concentration in the aqueous phase) for 10 min. Fixation was quenched by adding glycine to a final concentration of 250 mM and rotating for 5 min. Fixed embryos were then pelleted by centrifugation at 1,000 × *g* for 1 min and washed with PBST. Embryos were flash-frozen in liquid nitrogen and stored at -80 °C until use. For Micro-C, embryo fixation was performed following the previously described procedures^3,57^ with modifications. Embryos were fixed by rotating in a mixture of heptane and PBST containing 1.8% formaldehyde (Thermo Fisher; final concentration in the aqueous phase) for 10 min. Fixation was quenched by adding Tris-HCl (pH 7.5) to a final concentration of 0.94 M and rotation for 5 min. Embryos were pelleted by centrifugation at 1,000 × *g* for 1 min at 4 °C, washed with PBST, and then stored at 4 °C. At the end of the day, fixed embryos were pooled together and further fixed with PBST containing 3 mM DSG (Wako) for 45 min. Fixation was quenched by adding Tris-HCl (pH 7.5) to a final concentration of 0.54 M and rotation for 5 min. Embryos were pelleted by centrifugation at 1,000 × *g* for 1 min at 4 °C and then washed with PBST. Fixed embryos were flash-frozen in liquid nitrogen and stored at -80 °C until use.

### ChIP-seq

ChIP-seq was performed essentially as described in Saito *et al*.^57^ Approximately 50 µL of nc14 embryos were homogenized using a Dounce homogenizer in lysis buffer [15 mM HEPES (pH 7.5), 15 mM NaCl, 60 mM KCl, 4 mM MgCl2, 0.5% Triton X-100, 0.5 mM DTT, and 1× EDTA-free protease inhibitor (Nacalai Tesque)]. The nuclear suspension was sequentially washed twice with lysis buffer and once with wash buffer [15 mM HEPES (pH 7.5), 200 mM NaCl, 1 mM EDTA, 0.5 mM EGTA, and 1× EDTA-free protease inhibitor]. Nuclei were then resuspended in 300 µL of sonication buffer [15 mM HEPES (pH 7.5), 140 mM NaCl, 1 mM EDTA, 0.5 mM EGTA, 0.1% sodium deoxycholate, 0.5% sodium N-lauroylsarcosinate, and 1× EDTA-free protease inhibitor] and subjected to sonication using Bioruptor Pico (Diagenode) set to 30 sec ON / 30 sec OFF cycles, intensity HIGH, for 7 cycles. Four of the six available holders were used. Sonication was performed using 1.5 mL tubes (Diagenode). The resulting lysate was adjusted to a final volume of 600 µL containing 1% Triton X-100, and centrifuged at 20,000 × *g* for 10 min at 4 °C. The supernatant was collected, and one-twelfth of the sample (50 µL) was reserved as input and stored at -30 °C. We used 2 µg of anti-FLAG M2 monoclonal antibody (Sigma-Aldrich, F1804) conjugated with 10 µL of Dynabeads Protein G magnetic beads (Invitrogen) for immunoprecipitation (IP). IP was performed overnight at 4 °C. Beads were then washed five times with RIPA buffer [50 mM HEPES (pH 7.5), 500 mM LiCl, 1 mM EDTA, 1% NP-40, and 0.7% sodium deoxycholate] and once with TE buffer containing 50 mM NaCl. After centrifugation at 960 × *g* for 3 min at 4 °C, the supernatant was completely removed. Beads were resuspended in 210 µL of elution buffer [50 mM Tris-HCl (pH 8.0), 10 mM EDTA, and 1% SDS] and incubated at 65 °C for 30 min. The beads were collected using a magnetic stand and 200 µL of the supernatant was transferred to a new tube. For the input, 150 µL of elution buffer was added to the 50 µL reserved fraction. Reverse-crosslinking was performed by incubating both ChIP and input samples at 65 °C for 6-15 hours. Samples were treated with RNase A at 37 °C for 30 min and with proteinase K at 55 °C for an hour to remove RNA and proteins, respectively. DNA was purified by phenol-chloroform extraction followed by ethanol precipitation. The final DNA pellet was resuspended in 30 µL of water for the IP fraction or 50 µL for the input fraction. ChIP-seq libraries were prepared using the NEBNext Ultra II DNA Library Prep Kit for Illumina (NEB), starting with 25 µL of DNA. The adaptor was diluted 25-fold for ChIP DNA and 10-fold for input DNA. PCR was performed with 12 cycles for the IP fraction and 4 cycles for the input fraction. PCR products ranging from 100 to 1,000 bp were size-selected and purified using AMPure XP beads (Beckman Coulter). The quality of size-selected libraries was validated by TapeStation D5000HS kit (Agilent technology). The resulting libraries were sent to BGI Japan for circularization and sequenced on a DNBSEQ-G400 platform (MGI Tech) using paired-end 100 bp reads. Two biological replicates were prepared for each strain.

### Micro-C

Micro-C was performed essentially following the previously described procedures^3,57,106^ with some modifications. We used approximately 50 µL of dual-crosslinked embryos per replicate. Embryos were homogenized with Dounce homogenizer in 500 µL of MB1 buffer [10 mM Tris-HCl (pH 8.0), 50 mM NaCl, 5 mM MgCl2, 1 mM CaCl2, 0.2% NP-40, and 1× EDTA-free protease inhibitor], incubated on ice for 20 min, and washed once with MB1 buffer. Subsequently, chromatin was digested at 37 °C for 10 min in 100 µL of MB1 buffer with a pre-determined amount of Micrococcal Nuclease (NEB) to yield a ∼90% mononucleosome / ∼10% dinucleosome ratio. The digestion was terminated by adding 5 µL of 0.1 M EGTA and incubating at 65 °C for 10 min. After washing twice with MB2 buffer [50 mM NaCl, 10 mM Tris-HCl (pH 8.0), and 5 mM MgCl2], 45 µL of end-chewing mix [10 µL of 10 mM ATP, 2.5 µL of 100 mM DTT, 5 µL of 10× NEBuffer r2.1 (NEB), 25 µL of water, and 2.5 µL of 10 U/µL T4 Polynucleotide Kinase (NEB)] was added and incubated at 37 °C for 15 min. The mixture was then supplied with 5 µL of 5 U/µL Klenow Fragment (NEB) and incubated at 37 °C for 15 min. Subsequently, 25 µL of end-labeling mix [5 µL of 1 mM biotin-14-dATP (ActiveMotif), 5 µL of 1 mM biotin-14-dCTP (Jena Bioscience), 0.5 µL of 10 mM dGTP (Takara), 0.5 µL of 10 mM dTTP (Roche), 2.5 µL of 10× T4 DNA Ligase Buffer (NEB), 0.25 µL of 10 mg/mL BSA, and 11.25 µL of water] was added and incubated at 25 °C for 45 min for biotin labeling. The labeling was terminated by adding 5 µL of 0.5 M EDTA and incubating at 65 °C for 20 min. Chromatin was pelleted and washed with cold MB3 buffer [50 mM Tris-HCl (pH 8.0) and 10 mM MgCl2]. For proximal ligation, 250 µL of end-ligation mix [2.5 µL of 10 mg/mL BSA, 25 µL of 10× T4 DNA Ligase Buffer (NEB), 210 µL of water, and 12.5 µL of 400 U/µL T4 DNA Ligase (NEB)] was added and incubated for 4 h at room temperature with slow rotation. To remove biotin-dNTPs from unligated ends, 100 µL of unligated-end purification mix [10 µL of 10× NEBuffer 1 (NEB), 85 µL of water, and 5 µL of 100 U/µL Exonuclease III (NEB)] was added and incubated for 1 h at 37 °C. The sample was then reverse-crosslinked in elution buffer [370 mM NaCl, 46 mM Tris-HCl (pH 8.0), 9.3 mM EDTA, and 0.93% SDS] at 65 °C overnight. Samples were treated with RNase A at 37 °C for 30 min and with proteinase K at 55 °C for an hour to remove RNA and proteins, respectively. DNA was purified by phenol-chloroform extraction followed by ethanol precipitation. The final DNA pellet was resuspended in 20 µL of TE. Subsequently, the biotin-labeled DNA was captured by Dynabeads MyOne Streptavidin C1 beads (Invitrogen) in biotin binding buffer [5 mM Tris-HCl (pH 7.5), 0.5 mM EDTA, and 1 M NaCl] and incubated for 20 min at room temperature with rotation. Beads were then washed twice with tween wash buffer [5 mM Tris-HCl (pH 7.5), 0.5 mM EDTA, 1 M NaCl, and 0.05% Tween-20] and resuspended in 25 µL of 0.1× TE. The sequence libraries were prepared using the NEBNext Ultra II DNA Library Prep Kit for Illumina (NEB). End-repair and adaptor ligation were performed on beads. The adaptor was diluted 10-fold to minimize the formation of adaptor dimers. Beads were then washed with tween wash buffer and 0.1× TE, and libraries were amplified by PCR for 8 cycles. PCR products were purified using 0.9× AMPure XP beads (Beckman Coulter). The quality of libraries was validated by TapeStation D5000HS kit (Agilent technology). The resulting libraries were sent to BGI Japan for circularization and sequenced on a DNBSEQ-G400 platform (MGI Tech) using paired-end 100 bp reads. Two biological replicates were prepared for each strain.

### Live-imaging

Dechorionated embryos were mounted between a polyethylene membrane (Ube Film) and a coverslip (18 mm × 18 mm) and embedded in FL-100-450CS (Shin-Etsu Silicone). For Figure 2B and 2E, data acquisition was performed using Airyscan2 super-resolution imaging system with live embryos. Embryos were imaged using a Zeiss LSM 900 equipped with Airyscan2 detector. During imaging, temperature was kept in between 22.0 and 23.5°C. Plan-Apochromat 63× / 1.4 N.A. oil immersion objective was used. During imaging, data acquisition was occasionally stopped for a few seconds to correct z-position, and data were concatenated afterward. Raw z-stack images were subjected to 3D Airyscan processing using Zeiss Zen 3.1 software with the same parameter for the same set of experiments. Images were acquired with following settings: 940 × 940 pixels with pixel size 0.043 μm (40.05 × 40.05 μm regions), 16-bit depth, 89 z-slices separated by 0.17 μm at each time point (after Airyscan processing). For images in Figure 2B, fluorescences of CG31365-GFP-3×FLAG and His2Av-emiRFP670 were excited using 488-nm and 640-nm lasers, respectively, and every z-stack image was taken with a time resolution of 38.872 sec/frame. For images in Figure 2E, fluorescence of WT CG31365-GFP-3×FLAG or ΔZAD CG31365-GFP-3×FLAG was excited using 488-nm laser and every z-stack image was taken with a time resolution of 19.531 sec/frame. For Figure 1E, data acquisition was performed using a FV4000 confocal microscope equipped with SilVIR detector (Evident). UPlanXApo 40× / 1.4 N.A. oil immersion objective was used. Images were acquired with following settings: 1024 × 1024 pixels with pixel size 0.311 µm (318.198 µm × 318.198 µm regions), 16-bit depth, 16 z-slices separated by 1.0 µm. Fluorescence of His2Av-emiRFP670 was excited using 640-nm laser. Fly strains expressing His2Av-emiRFP670 were prepared by following procedures. For the control, first, *yw* adults were crossed with *His2Av-emiRFP670 (VK00033)* adults. Subsequently, resulting *His2Av-emiRFP670 (VK00033)/+* virgin females were mated with *yw* males. The resulting embryos were used for the imaging analysis. For the mutant, first, *CG31365[Δ1]/TM6* adults were crossed with *CG31365[Δ1], His2Av-emiRFP670 (VK00033)/TM6* adults. Subsequently, resulting *CG31365[Δ1]/CG31365[Δ1], His2Av-emiRFP670 (VK00033)* virgin females were mated with homozygous *CG31365[Δ1]* males from *CG31365[Δ1]/TM6* strain. The resulting embryos were used for the imaging analysis.

### Fluorescence recovery after photobleaching (FRAP)

Embryos were dechorionated in bleach, and then mounted between a polyethylene membrane (Ube Film) and a coverslip (18 mm × 18 mm) and embedded in FL-100-450CS (Shin-Etsu Silicone). Data acquisition was performed on a Zeiss LSM 900 microscope. During imaging, temperature was kept in between 22.0 and 23.5°C. Plan-Apochromat 63× / 1.4 N.A. oil immersion objective was used. Images were acquired using the following settings: 400 × 400 pixels with pixel size 0.063 μm (25.35 × 25.35 μm regions), 16-bit depth, 37 z-slices separated by 0.41 μm at each time point. Every z-stack image was taken with a time resolution of 5.906 sec/frame. Fluorescences of CG31365-GFP-3×FLAG and His2Av-emiRFP670 were excited using 488-nm and 640-nm lasers, respectively. Images were taken from the middle of nc13, and all the experiments were performed at ∼8-20 min into nc14. A circular ROI (0.5 µm × 0.5 µm) was bleached for 2.5-3.0 sec at the maximum laser intensity using “Interactive Bleaching” mode in Zeiss Zen 3.1 software. In total, 22 condensates from 13 embryos were analyzed. For some of the condensates, data acquisition was occasionally stopped for a few seconds to correct z-position during imaging, and data were concatenated afterward.

### Single-molecule inexpensive FISH and immunofluorescence staining

Single-molecule inexpensive FISH (smiFISH) was performed as previously described with some modifications.^107–109^ Embryos were collected over 30 min and aged for 2 h 10 min to reach at 2h10-2h40 after embryo laying. Collected embryos were dechorionated with bleach. Subsequently, embryos were shaken in fix solution buffer [0.5 mL of 10× PBS, 3.5 mL of water, 1 mL of 37% formaldehyde, and 5 mL of heptane] for 45 min at room temperature. After removing the aqueous solution layer, 10 ml of 100% methanol was added, and embryos were devitellinized by shaken for ∼1 min. Devitellinized embryos were washed with methanol and stored in methanol at -30°C before use. smiFISH probes targeting exons of endogenous *pyr*, *ths* (*ths-RA*) and *eve* genes were designed using the Biosearch Technologies Stellaris RNA FISH probe designer tool (https://www.biosearchtech.com/support/education/stellaris-rna-fish). The following sequence, which is the reverse complement of the X FLAP sequence described previously,^84^ was added to the 5′ end of each 20 nt probe: 5′-CCT CCT AAG TTT CGA GCT GGA CTC AGT G-3′. Oligos with the X FLAP sequence (5′-CAC TGA GTC CAG CTC GAA ACT TAG GAG G-3′) containing Cy3 or Cy5 both at the 5′ and 3′ ends were synthesized by Eurofins Genomics. Probes and the X FLAP were annealed and then stored at -30°C before use. Probes for *pyr* and *ths* were annealed to Cy3 oligos, and probes for *eve* and MS2 were annealed to Cy5 oligos. The sequences of smiFISH probes used in this study are listed in Table S1. Hybridization was performed with some modifications from the previously described procedures.^107–109^ Fixed embryos stored in 100% methanol were transferred to PBST [1× PBS and 0.05% Tween 20], and washed with PBST and smiFISH wash buffer [2× SSC and 10% deionized formamide]. Subsequently, embryos were incubated in smiFISH wash buffer at 37°C for 2× 30 min. Then, annealed smiFISH probes diluted in smiFISH hybridization buffer [10% w/v dextran sodium sulphate 5000, 2× SSC, and 10% deionized formamide] to a final concentration of 80 nM were added, and hybridized with embryos in the dark at 37°C for over 14 h. Embryos were then washed 4× 15 min in pre-warmed smiFISH wash buffer at 37°C, and 3× 10 min in PBST at room temperature. For images in Figure 5B, embryos were stained with DAPI solution for 10 min, then washed with PBST. Embryos were then mounted in ProLong Gold Antifade Mountant (Thermo Fisher) for imaging. For images in Figure 6A-D, in which *CG31365-GFP-3×FLAG* strain was used for the analysis, immunofluorescence staining using anti-FLAG antibody was performed following the hybridization step, with some modifications from the previously described procedures.^107–109^ Embryos were blocked for 1 h in blocking solution [1.5× western blocking reagent (Roche) in PBST], and then incubated with mouse anti-FLAG M2 monoclonal antibody (Sigma-Aldrich, F1804) diluted 1:500 in blocking solution for 1 h at 25 °C. Embryos were then washed with PBST, blocked for 1 h in blocking solution, and incubated with secondary antibody (anti-mouse IgG Alexa Fluor 488, Thermo Fisher, A-21202) diluted 1:500 in blocking solution for 2 h at room temperature. After washing with PBST, embryos were stained with DAPI solution for 10 min, then washed with PBST. Lastly, embryos were mounted in ProLong Gold Antifade Mountant (Thermo Fisher) for imaging.

### Imaging of smiFISH and immunofluorescence staining embryos

For images in Figure 5B, data acquisition was performed using a FV4000 confocal microscope equipped with SilVIR detector (Evident). UPlanXApo 40× / 1.4 N.A. oil immersion objective was used. Images were acquired with following settings: 2048 × 2048 pixels with pixel size 0.155 µm (318.198 µm × 318.198 µm regions), 16-bit depth, 10 z-slices separated by 1.1 µm. Fluorescences of DAPI, Cy3, and Cy5 were excited using 405-nm, 561-nm and 640-nm lasers, respectively. For images in Figure 6A-D, data acquisition was performed using Airyscan2 super-resolution imaging system. Embryos were imaged using a Zeiss LSM 900 equipped with Airyscan2 detector. Plan-Apochromat 63× / 1.4 N.A. oil immersion objective was used. Raw z-stack images were subjected to 3D Airyscan processing using Zeiss Zen 3.1 software with the same parameter for the same set of experiments. Images were acquired with following settings: 2186 × 2186 pixels with pixel size 0.035 μm (77.16 × 77.16 μm regions), 16-bit depth, 58 z-slices separated by 0.14 μm (after Airyscan processing). Fluorescence of DAPI, Alexa488, Cy3, and Cy5 were excited using 405-nm, 488-nm, 561-nm and 640-nm lasers, respectively. Before final image acquisition, the regions showing overlapped expression of *pyr/ths* and *eve* were identified by acquiring tiled images. Based on wide-field views and DAPI signals, we consistently acquired images from early nc14 embryos.

### Identification and analysis of CG31365 orthologs

Using the amino acid sequence of *D. melanogaster* CG31365 as a query, we performed a blastp search (https://blast.ncbi.nlm.nih.gov/Blast.cgi). Target sequences were filtered to retain only those annotated in the latest genome assemblies and gene annotations listed below: dm6 (FlyBase Release 6.54) for *D. melanogaster*, ASM438219v2 (NCBI Release 101) for *D. sechellia*, Prin_Dsim_3.1 (NCBI Release 103) for *D. simulans*, DereRS2 (NCBI Release 101) for *D. erecta*, Prin_Dyak_Tai18E2_2.1 (NCBI Release 102) for *D. yakuba*, ASM1815383v1 (NCBI Release 102) for *D. eugracilis*, RU_DBia_V1.1 (NCBI Release 103) for *D. biarmipes*, CBGP_Dsuzu_IsoJpt1.0 (GCF_043229965.1-RS_2025_01) for *D. suzukii*, DtakHiC1v2 (GCF_030179915.1-RS_2024_12) for *D. takahashii*, ASM1815250v1 (NCBI Release 102) for *D. elegans*, ASM1815211v1 (NCBI Release 102) for *D. rhopaloa*, ASM1815226v1 (NCBI Release 102) for *D. ficusphila*, DkikHiC1v2 (GCF_030179895.1-RS_2024_12) for *D. kikkawai*, ASM1763931v2 (NCBI Release 102) for *D. ananassae*, DperRS2 (NCBI Release 101) for *D. persimilis*, UCI_Dpse_MV25 (GCF_009870125.1-RS_2025_01) for *D. pseudoobscura*, UCI_dwil_1.1 (NCBI Release 102) for *D. willistoni*, ASM1815329v1 (NCBI Release 103) for *D. grimshawi*, ASM1815372v1 (NCBI Release 102) for *D. mojavensis*, Dvir_AGI_RSII-ME (GCF_030788295.1-RS_2024_12) for *D. virilis* and AaegL5.0 (NCBI Release 101) for *A. aegypti*. Then, for each species, the sequence with the highest percentage identity was defined as the ortholog. Orthologous sequences were aligned using CLUSTALW (https://www.genome.jp/tools-bin/clustalw), and a molecular phylogenetic tree was constructed with the FastTree method.^110^ For Figure S1A, sequence similarity between each ortholog and *D. melanogaster* CG31365 was visualized on the pairwise alignment using the BLOSUM62 matrix (https://www.ncbi.nlm.nih.gov/IEB/ToolBox/C_DOC/lxr/source/data/BLOSUM62) as a proxy. The phylogenetic tree was visualized using ETE toolkit.^100^ The amino acid sequences aligned to the N-terminal ZAD in *D. melanogaster* CG31365 were extracted and used for structural prediction analysis.

### Structural prediction and visualization

Structures of CG31365 ZAD homodimers were predicted using AlphaFold3^77^ and visualized using ChimeraX.^78^ For Figure S1B, to visualize orthologous ZAD dimers, the matchmaker command was used to superimpose each predicted structure onto the reference structure, *i.e.*, the predicted structure of *D. melanogaster* CG31365 ZAD homodimer. The reference structure was removed prior to image generation.

### Plasmids

#### gRNA expression plasmids

Two DNA oligos containing gRNA sequences were annealed and inserted into the pCFD3-dU6:3gRNA plasmid (addgene # 49410) using BbsI sites. Sequences of DNA oligos used for gRNA cloning are listed in Table S2.

#### pBS-CG31365[Δ1] 5′ arm-loxP-3×P3-dsRed-loxP-CG31365[Δ1] 3′ arm

A DNA fragment containing 5′ homology arm upstream of *CG31365* ORF was PCR amplified from *yw* genomic DNA and digested with SalI. The resulting fragment was inserted at the SalI site in the pBS-loxP-dsRed-loxP plasmid.^109^ Subsequently, a DNA fragment containing 3′ homology arm downstream of *CG31365* ORF was PCR amplified from *yw* genomic DNA and digested with NotI. The resulting fragment was inserted at the NotI site in the plasmid containing the 5′ homology arm.

#### pBS-CG31365 5′ arm-GFP-3×FLAG-loxP-3×P3-dsRed-loxP-CG31365 3′ arm

A DNA fragment containing 5′ homology arm of *CG31365* was PCR amplified from *yw* genomic DNA and digested with KpnI and SalI. The resulting fragment was inserted between the KpnI site and the SalI site in the pBS-GFP-3×FLAG-loxP-3×P3-dsRed-loxP plasmid.^108^ Subsequently, a DNA fragment containing 3′ homology arm of *CG31365* was PCR amplified from *yw* genomic DNA and digested with SpeI and NotI. The resulting fragment was inserted between the SpeI site and the NotI site in the plasmid containing the 5′ homology arm.

#### pBS-Mulb site 5′ arm-loxP-3×P3-GFP-loxP-Mulb site 3′ arm

A DNA fragment containing 3′ homology arm downstream of the CG31365/Mulberry enrichment site at *pyr* promoter was PCR amplified from *yw* genomic DNA and digested with NotI. The resulting fragment was inserted at the NotI site in the pBS-loxP-GFP-loxP plasmid.^111^ Subsequently, a DNA fragment containing 5′ homology arm upstream of the CG31365/Mulberry enrichment site at *pyr* promoter was PCR amplified from *yw* genomic DNA and digested with KpnI and SalI. The resulting fragment was inserted between the KpnI site and the SalI site in the plasmid containing the 3′ homology arm.

#### pBS-Enhancer 5′ arm-loxP-3×P3-GFP-loxP-Enhancer 3′ arm

A DNA fragment containing 5′ homology arm upstream of the shared enhaner at *pyr/ths* locus was PCR amplified from *yw* genomic DNA and digested with KpnI and SalI. The resulting fragment was inserted between the KpnI site and the SalI site in the pBS-loxP-GFP-loxP plasmid.^111^ Subsequently, a DNA fragment containing 3′ homology arm downstream the shared enhaner was PCR amplified from *yw* genomic DNA and digested with SpeI and NotI. The resulting fragment was inserted between the SpeI site and the NotI site in the plasmid containing the 5′ homology arm.

#### pBS-ths TSS 5′ arm-loxP-3×P3-GFP-loxP- ths TSS 3′ arm

A DNA fragment containing 5′ homology arm upstream of the *ths-RA* TSS was PCR amplified from *yw* genomic DNA and digested with KpnI and SalI. The resulting fragment was inserted between the KpnI site and the SalI site in the pBS-loxP-GFP-loxP plasmid.^111^ Subsequently, a DNA fragment containing 3′ homology arm downstream the *ths-RA* TSS was PCR amplified from *yw* genomic DNA and digested with SpeI and NotI. The resulting fragment was inserted between the SpeI site and the NotI site in the plasmid containing the 5′ homology arm.

#### pBS-CG31365 CDS^WT^

A DNA fragment containing *CG31365* CDS was PCR amplified from *yw* cDNA using primers (5′-AAA AGG TAC CAT GGA TGC ACA GCC GTC GTA TCC-3′) and (5′- AAA AGG TAC CGG CCA CAG CTT GAA ACT CGT TCG T-3′) and digested with KpnI. The resulting fragment was inserted at the KpnI site in the pBlueScript SK II (-) plasmid.

#### pBS-CG31365 CDS^ΔZAD^

Coding sequence for ZAD domain was specifically removed from pBS-CG31365 CDS^WT^ plasmid by PCR-based site-directed mutagenesis using primers (5′-CGT CGT ATC CCG ATT TGG CGA TCC GAC TTC TGG GTC TCG G-3′) and (5′-CCG AGA CCC AGA AGT CGG ATC GCC AAA TCG GGA TAC GAC G-3′).

#### pbφ-GFP-3×FLAG

A DNA fragment containing GFP-3×FLAG sequence was PCR amplified from pBS-GFP-3×FLAG-loxP-3×P3-dsRed-loxP plasmid^108^ using primers (5′-AAA ACT CGA GAA GCT TGG TAC CGG CGG CTC GGG TTC GGG ATC-3′) and (5′-AAA AGG ATC CTT ACT TGT CGT CAT CAT CCT TGT AGT CAA-3′) and digested with XhoI and BamHI. The resulting fragment was inserted between the XhoI and BamHI sites in the pbφ-multi cloning site plasmid.^112^

#### pbφ-GFP-3×FLAG-CG31365 3′ region

A DNA fragment containing 3′ region (3′ UTR and downstream ∼500 bp) of *CG31365* was PCR amplified from *yw* genomic DNA and digested with BamHI and XbaI. The resulting fragment was inserted between the BamHI and XbaI sites in the pbφ-GFP-3×FLAG plasmid.

#### pbφ-CG31365 5′ region-GFP-3×FLAG-CG31365 3′ region

A DNA fragment containing 5′ region (5′ UTR and upstream ∼2 kbp) of *CG31365* was PCR amplified from *yw* genomic DNA and digested with NotI and KpnI. The resulting fragment was inserted between the NotI and KpnI sites in the pBlueScript SK II (-) plasmid. This fragment contains the entire ORF of *CG31457* gene. To avoid cloning intact ORF, PCR-based site-directed mutagenesis was performed using primers (5′-TTG CCT TGA AAT TCA ACA TGA TGT TTG TCG CCT GTG CTT G-3′) and (5′-CAA GCA CAG GCG ACA AAC ATC ATG TTG AAT TTC AAG GCA A-3′) to induce a single-nucleotide deletion just downstream of the start codon of *CG31457*. The resulting plasmid was digested with NotI and KpnI, and the resulting DNA fragment containing the mutagenized 5′ region was inserted between the NotI and KpnI sites in the pbφ-GFP-3×FLAG-CG31365 3′ region plasmid.

#### pbφ-CG31365 5′ region-CG31365 CDS^WT^-GFP-3×FLAG-CG31365 3′ region

pBS-CG31365 CDS^WT^ was digested with KpnI, and the resulting DNA fragment containing *CG31365* CDS^WT^ was inserted at the KpnI site in the pbφ-CG31365 5′ region-GFP-3×FLAG-CG31365 3′ region plasmid.

#### pbφ-CG31365 5′ region-CG31365 CDS^ΔZAD^-GFP-3×FLAG-CG31365 3′ region

pBS-CG31365 CDS^ΔZAD^ was digested with KpnI, and the resulting DNA fragment containing *CG31365* CDS^ΔZAD^ was inserted at the KpnI site in the pbφ-CG31365 5′ region-GFP-3×FLAG-CG31365 3′ region plasmid.

#### pbφ-Mulb site-DSCP-24×MS2-yellow

A DNA fragment containing Mulb site (the CG31365/Mulberry enrichment site at *pyr* promoter, corresponding to the genomic region deleted in our *ΔMulb site* strain) was PCR amplified from *yw* genomic DNA and digested between NotI and XhoI. The resulting fragment was inserted between NotI site and XhoI site in pbφ-DSCP-MS2-yellow plasmid.^113^

#### pbφ-Enhancer-DSCP-24×MS2-yellow

A DNA fragment containing Enhancer (the shared enhaner at *pyr/ths* locus, corresponding to the genomic region deleted in our *ΔEnhancer* strain) was PCR amplified from *yw* genomic DNA and digested between NotI and XhoI. The resulting fragment was inserted between NotI site and XhoI site in pbφ-DSCP-MS2-yellow plasmid.^113^

### Quantification and statistical analysis Computational analysis of public RNA-seq data

Stage-specific RNA-seq data from *yw* nc14 embryos^36^ (ArrayExpress E-MTAB-11978; ENA run accession numbers ERR9951951, ERR9951952, ERR9951953, ERR9951954, ERR9951955, ERR9951956, ERR9951957, ERR9951958, ERR9951959) was used to comparatively examine the expression levels of ZAD-ZnF genes. Low-quality reads were removed and adapter sequences were trimmed from raw reads using fastp^85^ with -c option. The trimmed reads were first mapped to dm6 ncRNAs (BDGP6.32) using STAR^86^ with - -outFilterMultimapNmax 2000 --outReadsUnmapped Fastx options to retain unaligned reads. The unaligned reads were then mapped to dm6 genome (BDGP6.32) using STAR.^86^ Since the results showed high correlations among biological replicates, samtools^87^ merge command was used to merge the BAM files for each replicate. Subsequently, transcript abundance was quantified in TPM using StringTie^88^ with -e option based on dm6 annotations (BDGP6.32). For Figure 1A, the results for previously annotated ZAD-ZnF genes^54^ were filtered and visualized.

### Computational analysis of ChIP-seq data

Low-quality reads were removed and adapter sequences were trimmed from raw reads using fastp.^85^ The trimmed reads were then mapped to dm6 genome (BDGP6.32) with Bowtie2.^89^ Only uniquely mapped reads were retained using samtools^87^ view command with -F 0x4 -q 42 options. The resulting BAM files were sorted and indexed using samtools^87^ sort and index commands, respectively. Since the results showed high correlations among biological replicates, samtools^87^ merge command was used to merge the BAM files for each replicate. We used this merged dataset for all subsequent analyses. Peak calling was performed using MACS2^90^ with the FLAG-ChIP data for *yw* strain (a parental strain for all genome-edited and transgenic strains) as a control with -f BAMPE -g dm -q 0.01 options. For all visualizations of ChIP-seq profiles obtained in this study, CPM-normalized ChIP tracks with input-subtracted signals were used. These BigWig tracks were generated from BAM files using bamCompare command provided by deepTools.^91^ For Figure 3B, input-subtracted ChIP-seq profiles were converted to BedGraph format using UCSC bigWigToBedGraph.^93^ For Figure 3C and S2D, 499 bp-long genomic regions were defined around each peak summit using bedTools,^92^ and these regions were subjected to motif analysis using MEME^80^ provided in MEME Suite.^114^ For Figure 3D, 4C, S2B, and S2E, heatmap was plotted using computeMatrix and plotHeatmap commands provided by deepTools.^91^ For Figure 3D, S2B, and S2E, ChIP-seq tracks from a previous publication^36^ were used. For Figure 4A, 4B, and S4A, visualization of ChIP-seq tracks was performed using Integrative Genome Viewer (IGV).^81^ For Figure 5A and S4A, ChIP-seq tracks from previous publications (Zld in 120–150 min AEL embryos [GEO accession number GSM763061^82^] and Dl in 2-4 h AEL embryos [GEO accession number GSM1341814^83^]) were used. In addition, for Figure S4A, ChIP-seq track from previous publication (Pol II in 2h10-2h40 AEL embryos [GEO accession numbers GSM8951782 and GSM8951783^57^]) was used. When necessary, downloaded BigWig files were converted to BedGraph files using UCSC bigWigToBedGraph^93^ and lifted from dm3 to dm6 using UCSC liftOver.^94^ For Figure S2C, peaks with summits overlapping with +/- 250 bp around transcription start sites (TSS) or transcription end sites (TES) were defined to be located at TSS or TES, respectively. For the remaining peaks, those with summits overlapping with gene bodies spanning ≥500 bp were assigned to intragenic regions, while all other peaks were assigned to intergenic regions. The pie chart was generated in R based on dm6 annotations (BDGP6.32). Quality metrics for ChIP-seq libraries generated in this study are summarized in Table S3.

### Computational analysis of Micro-C data

Micro-C data was processed essentially as described in Batut *et al*.^3^ with some modifications. Low-quality reads were removed and adapter sequences were trimmed from raw reads using fastp.^85^ The trimmed reads were mapped to dm6 genome (BDGP6.32) using BWA^95^ with mem -SP5M options. The mapped reads were parsed with a filter to retain reads with a mapping quality ≥ 3, followed by sorting and deduplication, using pairtools.^96^ Subsequently, reads were selected with ’(pair_type == “UU”) or (pair_type == “UR”) or (pair_type == “RU”)’ option using pairtools.^96^ Matrix aggregation and balancing were performed using Cooler.^97^ Visual inspection of contact matrices by HiGlass^98^ showed strong concordance among biological replicates. Accordingly, the biological replicates were merged after deduplication to generate final high-coverage contact matrices for each genotype. We also used Higlass^98^ to manually inspect and compare differential contact profiles between the control and mutant strains. For visualization of Micro-C contact-frequency maps, subtraction heatmaps and virtual 4C profiles, CoolBox toolkit^79^ was used. For these maps, 400-bp resolution contact matrices were used, and balanced contact counts were log-10 transformed. The default “germany” colormap from FAN-C^99^ visualization was adopted in contact-frequency maps for improved readability. For virtual 4C profiles, previously annotated loop anchors^3^ (GSE171396) were normalized to 2 kbp and used as viewpoints. For Figure 3B, insulation scores were calculated from *yw* contact matrix using FAN-C^99^ with -w 10kb option, and the resulting file was converted to BED format. Quality metrics for Micro-C libraries generated in this study are summarized in Table S4.

### Image analysis for FRAP experiment

All z-stack images were maximum projected using Fiji.^101^ Then, for each image, a 60 × 60 pixel-size area (3.80 × 3.80 μm) around the photobleached condensate was manually determined for signal detection. We used condensates that stayed within these 60 × 60 pixel-size areas for the subsequent analysis, as most condensates remained largely stable in position during experiments (∼200 sec) in the developmental stage at which FRAP experiments were performed (∼8-20 min into nc14; see Video S1). Images were then processed for condensate intensity analysis using MATLAB. For each timepoint, images were median filtered using 3 × 3 kernels. Pixels with the lowest intensity in the median-filtered images were determined, and their corresponding intensity values were extracted from the raw images as background signal values. Subsequently, the pixels with the highest intensity in the median-filtered images were identified within the predetermined 60 × 60 pixel-size area. Then, using the 11 × 11 pixel-sized areas around these pixels (0.70 × 0.70 µm), the average intensity values were measured. The condensate signal values were calculated by subtracting the background signal values from the average intensity values. After calculating the condensate signal values from every timepoint, the signal values were smoothed using a moving average over 3 time points. The signal values were then normalized to the recovery rates after photobleaching (the average signal values before photobleaching = 1, the signal value immediately after photobleaching = 0). In total, 22 condensates from 13 embryos were analyzed. For the representative images of photobleached condensate in Figure 2C, the 15 × 15 pixel-size area (0.95 × 0.95 µm) around the pixel with the highest intensity in the median-filtered image was cropped from the raw z-projected image. To improve readability, the cropped images were then median filtered using Fiji^101^ with a radius of 1.

### Image analysis for smiFISH and immunofluorescence experiment

First, images were preprocessed using Fiji^101^ as follows. To avoid detecting false-positive DAPI signals from yolk regions, 40 z-stacks around DAPI-stained nuclei were cropped from raw z-stack images for each image. Using these cropped images, 3D nuclear masks were computed from DAPI images based on the threshold values automatically determined by the Otsu method using Gaussian-blurred maximum projected images. Next, images were processed for signal analysis using MATLAB. RNA FISH spots were detected using Airlocalize.^102^ Parameters for Airlocalize software were interactively optimized for each image to minimize false-positive detection. Subsequently, for each image, single-molecule RNA intensity values were determined as the mode value in the histogram intensity profile. Using this single-molecule spot intensity, signals of all the detected FISH spots were normalized to absolute number of RNA molecules. Considering the differences in expression levels among genes, threshold intensity values to define active transcription sites were determined as follows: 3 RNA molecules for *pyr* and *ths* genes, and 8 RNA molecules for *eve* gene. Then, spots with molecules above these thresholds and involved in the predetermined 3D nuclear mask were defined as active transcription sites. In total, 5 embryos were analyzed for the *pyr* and *eve* co-staining experiment, whereas 6 embryos were analyzed for the *ths* and *eve* co-staining experiment. For the average profile plots in Figure 6C and 6D, the 31 × 31 × 3 pixel-size areas around the active transcription sites (1.09 × 1.09 × 0.42 µm) were used. Intensity values in each pixel position were averaged across all the active transcription sites, and z-stack profiles were then average projected.

### Statistical analysis

For all statistical analyses, the tests used, as well as specifics about the data sets to which these tests were applied, are specified in the appropriate figure legend.

