## Supplementary material for "The *Drosophila* ZAD zinc finger protein Mulberry shapes the organization of the regulatory genome in the early embryo": Figure S1-7

### Umemura *et al.*, Figure S1

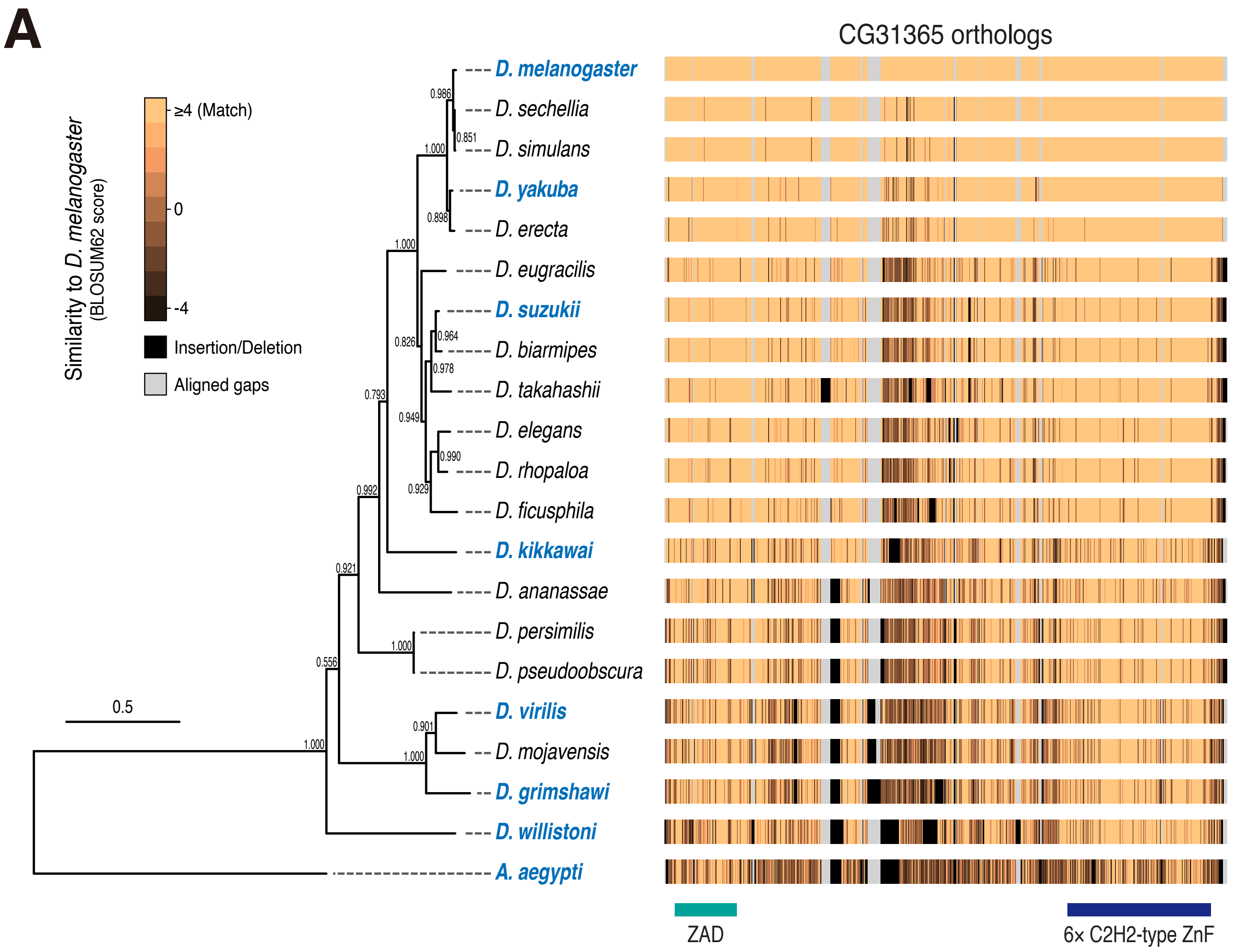


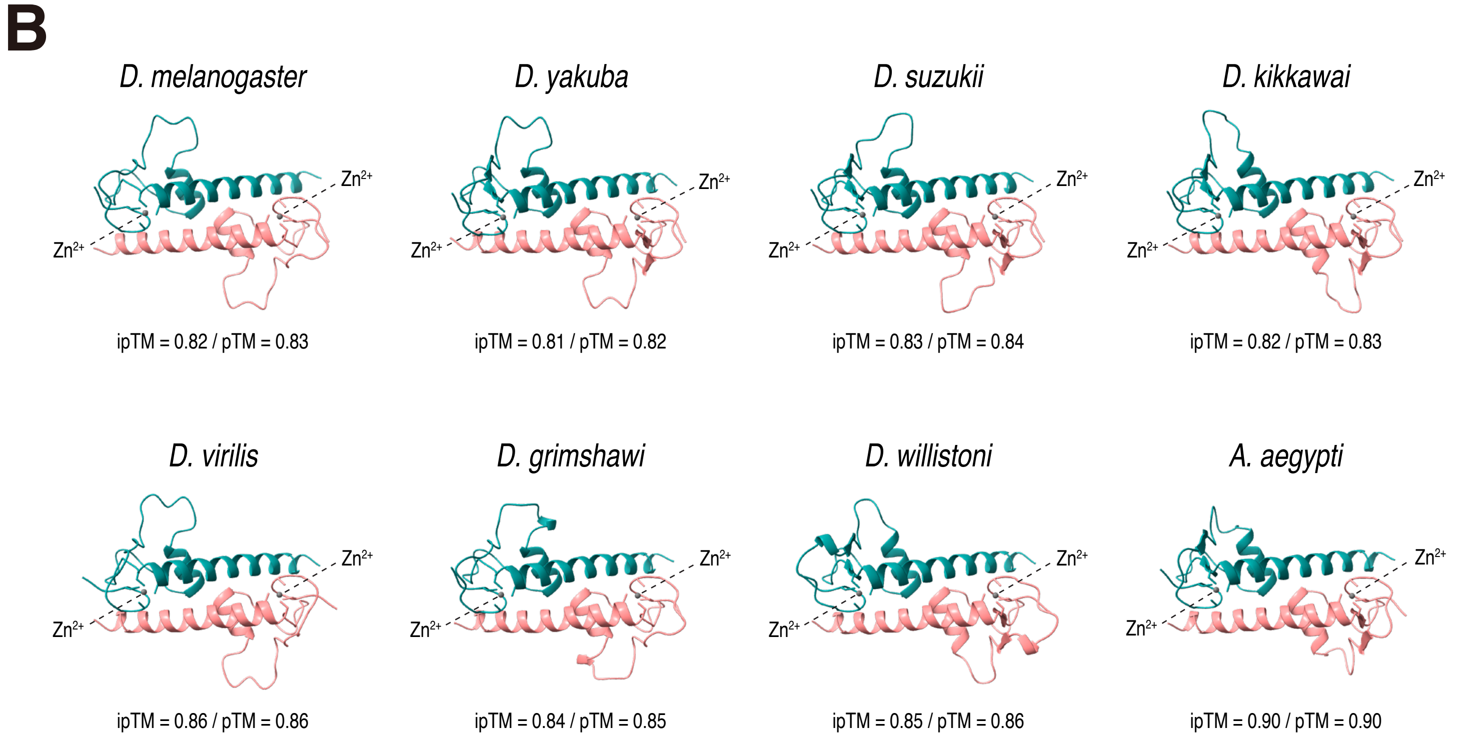

**Figure S1. Evolutionary conservation of CG31365 ZAD, related to Figure 2**

(A) Molecular phylogenetic tree alongside pairwise alignment of CG31365 orthologs. The tree was rooted using the *A. aegypti* ortholog as an outgroup. On the alignment, sequence similarity to *D. melanogaster* CG31365 is visualized. The names of species for orthologs used in structural prediction of ZAD dimers are shown in blue.

(B) AlphaFold3 prediction of the ZAD dimers for representative CG31365 orthologs. Protein structures were visualized using UCSF ChimeraX.[78] For *D. melanogaster*, the data is the same as that shown in Figure 2D.

Umemura *et al.*, Figure S2

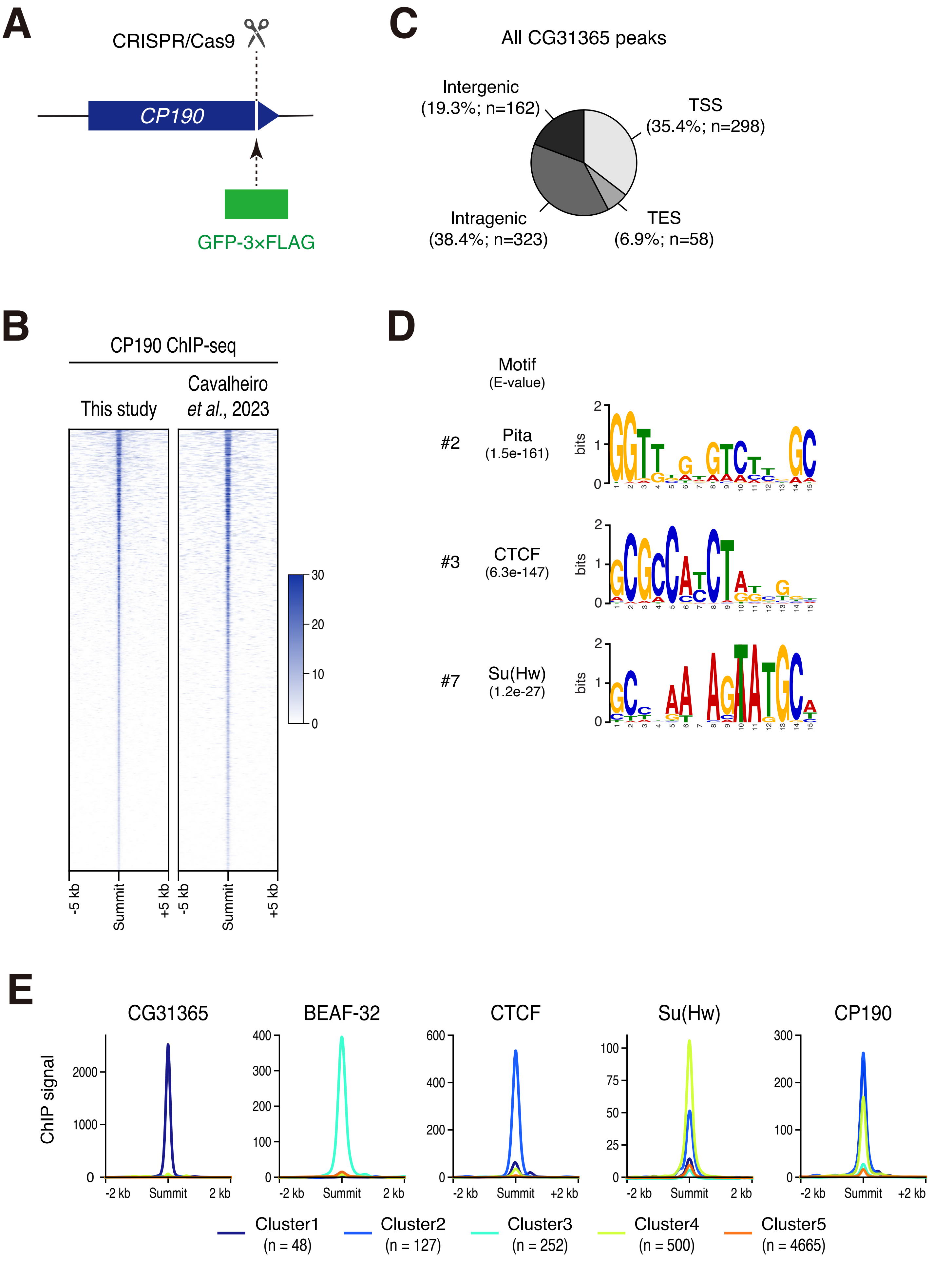

**Figure S2. Details about ChIP-seq experiments, related to Figure 3**

(A) Schematic representation of CRISPR/Cas9-mediated GFP-3×FLAG tagging for CP190 protein, previously performed in Saito *et al.*[57]

(B) Heatmap visualization of CP190 ChIP-seq profile in our analysis using anti-FLAG antibody (left) and previously reported ChIP-seq profile using anti-CP190 antibody (right).[36] Profiles are visualized at CP190 peaks identified in our analysis using anti-FLAG antibody and sorted by the signal intensity.

(C) Pie chart showing distribution profiles of endogenous CG31365 ChIP-seq peaks.

(D) The motif sequence logo of Pita, CTCF, and Su(Hw) at CG31365 peaks, identified by MEME.[80] The relative ranking of enrichment among all the significantly enriched motifs is indicated in the figure.

(E) Averaged ChIP-seq profiles at CP190 peaks, clustered by k-means (n=5). The clustering result is the same as in Figure 3D. Clusters 1–4 were named sequentially from top to bottom based on the heatmap in Figure 3D. Cluster 5 is omitted in Figure 3D, because it showed considerably small enrichment across all ChIP-seq profiles. The counts of analyzed profiles in each cluster are indicated in the figure. For BEAF-32, CTCF, and Su(Hw), publicly available ChIP-seq tracks in nc14 embryos[36] were used for the visualization.

**A**

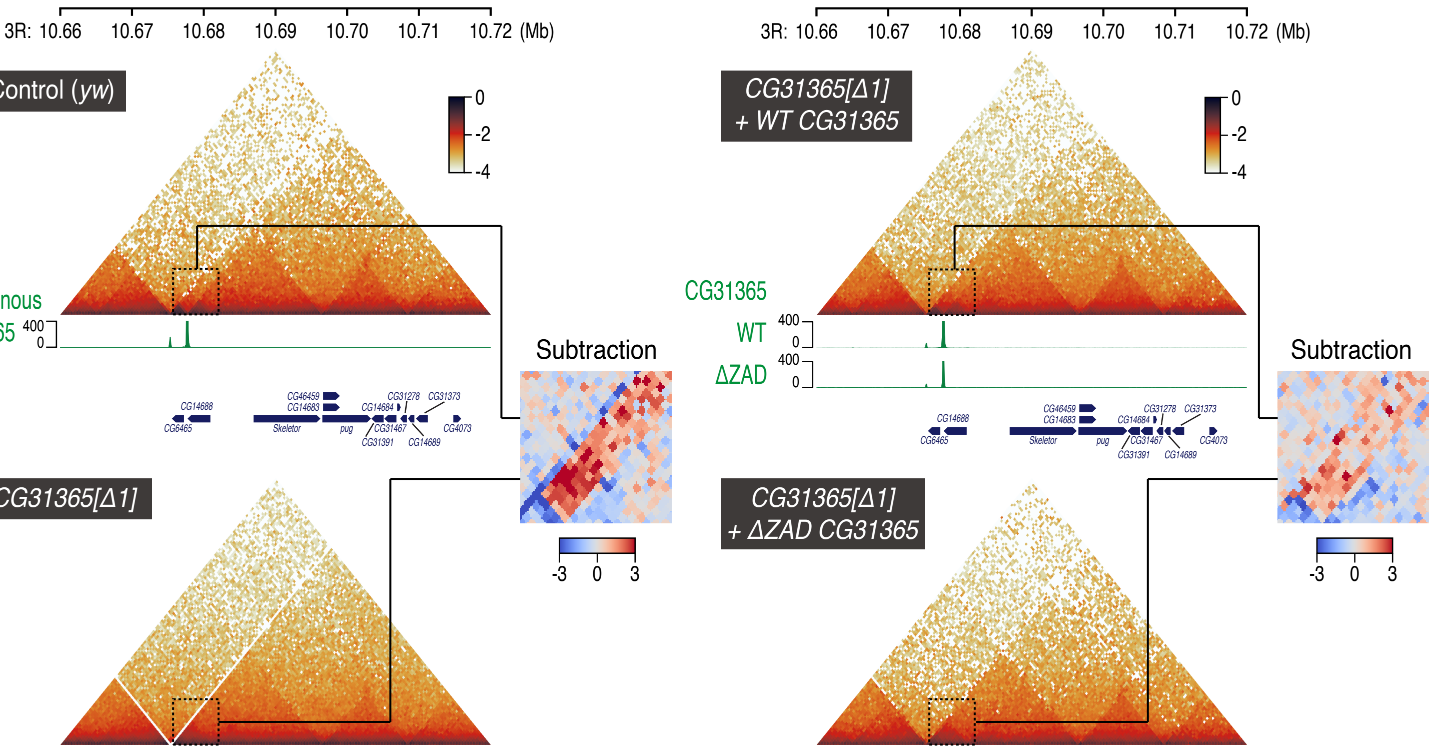
3R: 10.66 10.67 10.68 10.69 10.70 10.71 10.72 (Mb)
Control (yw)
0
-2
-4
Endogenous
CG31365
400
0
CG6465
CG14688
Skeletor
pug
CG46459
CG14683
CG14684
CG31391
CG31467
CG31278
CG31373
CG14689
CG4073
Subtraction
-3 0 3
CG31365[Δ1]
CG31365[Δ1]
+ WT CG31365
CG31365
WT
ΔZAD
CG31365[Δ1]
+ ΔZAD CG31365

**B**

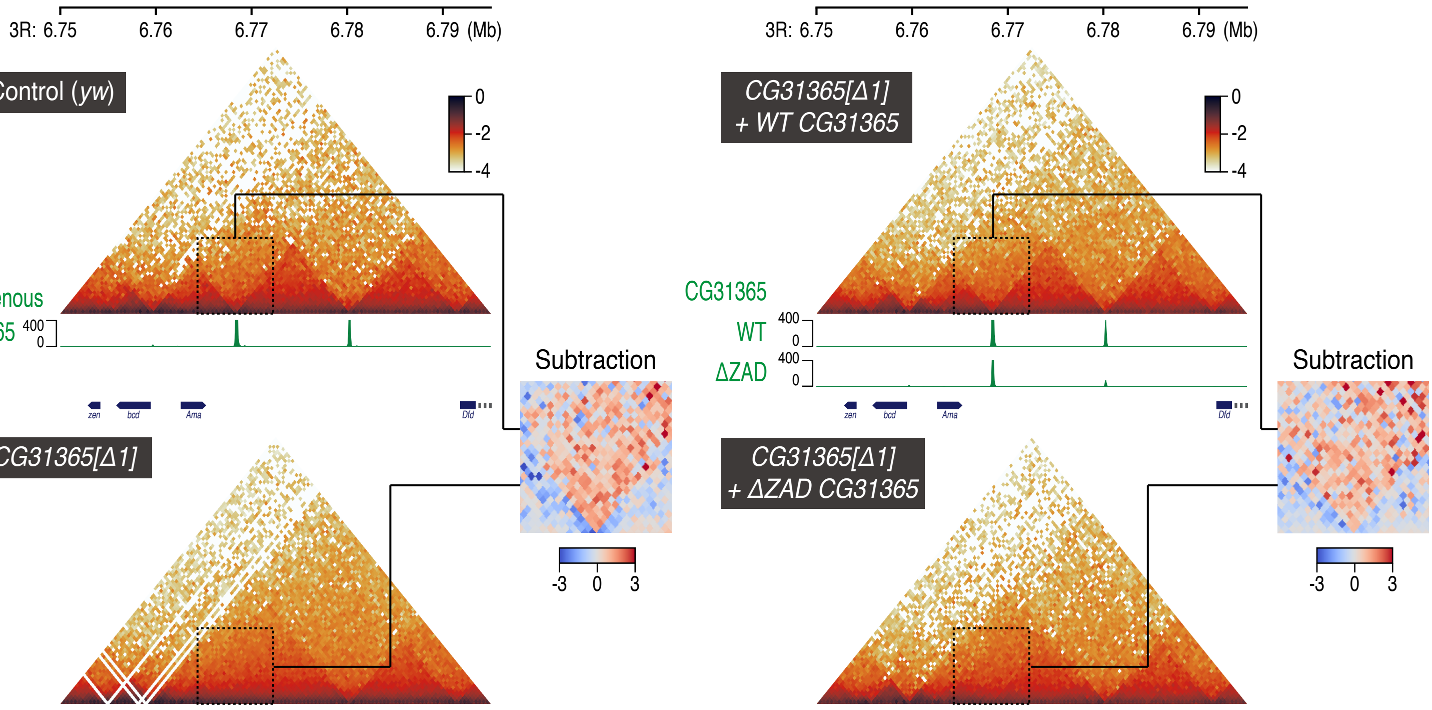
3R: 6.75 6.76 6.77 6.78 6.79 (Mb)
Control (yw)
0
-2
-4
Endogenous
CG31365
400
0
zen
bcd
Ama
Dfd
Subtraction
-3 0 3
CG31365[Δ1]
CG31365[Δ1]
+ WT CG31365
CG31365
WT
ΔZAD
CG31365[Δ1]
+ ΔZAD CG31365

**Figure S3. Topological boundaries regulated by CG31365, related to Figure 4**

(A, B) Micro-C contact-frequency maps are shown alongside ChIP-seq tracks. Micro-C experiments were performed using corresponding homozygous strains. ChIP-seq experiments were also performed using corresponding homozygous strains, except that *CG31365-GFP-3×FLAG* strain was used for endogenous CG31365. Differential contact profiles are summarized in subtraction heatmaps (*Mulb[Δ1]* − *yw* and *ΔZAD CG31365* rescue − *WT CG31365* rescue). Coolbox toolkit[79] was used for visualization.

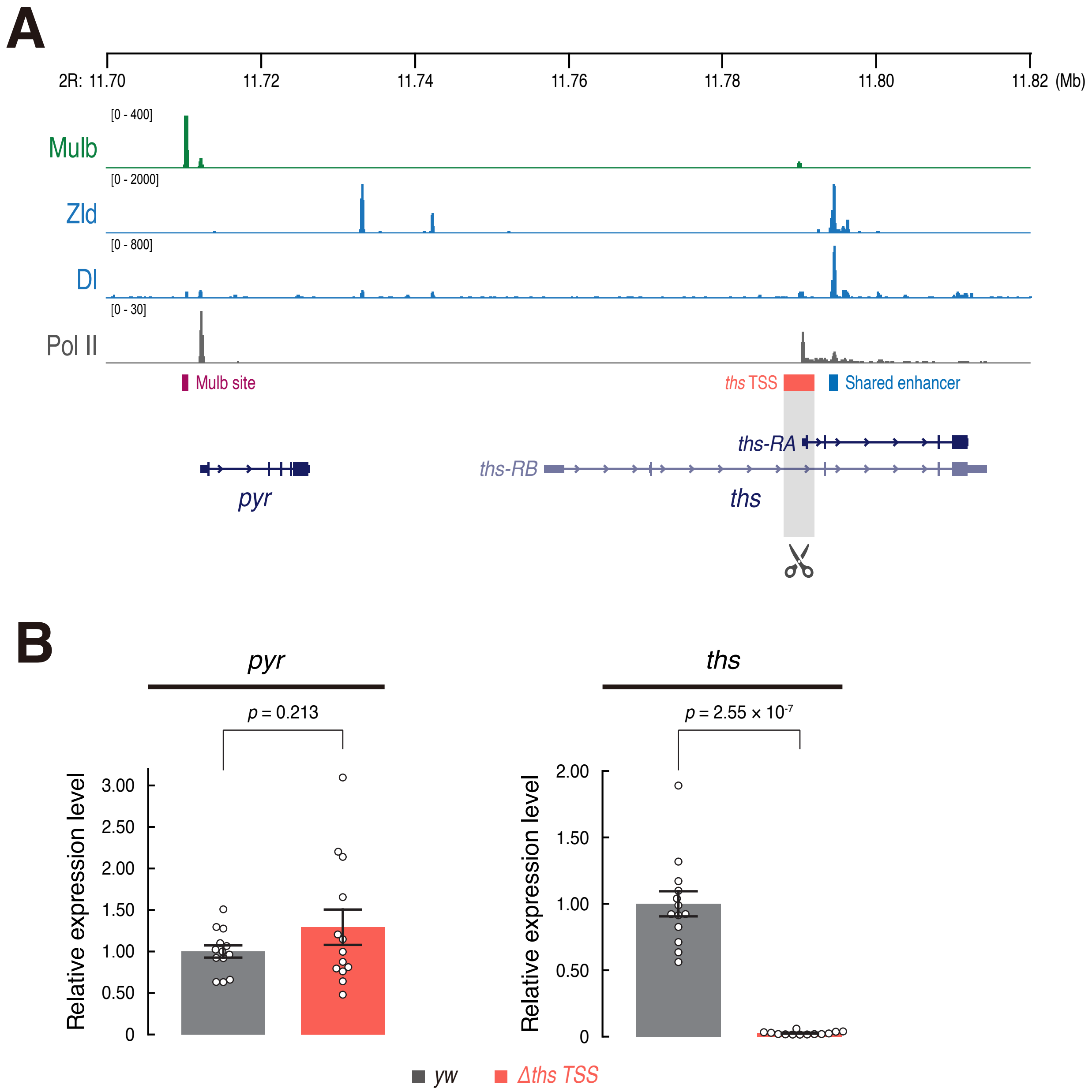
A
2R: 11.70
11.72
11.74
11.76
11.78
11.80
11.82 (Mb)
[0 - 400]
Mulb
[0 - 2000]
Zld
[0 - 800]
Dl
[0 - 30]
Pol II
Mulb site
ths TSS
Shared enhancer
ths-RA
ths-RB
pyr
ths
B
pyr
ths
p = 0.213
p = 2.55 × 10-7
Relative expression level
3.00
2.50
2.00
1.50
1.00
0.50
0
yw
Δths TSS

**Figure S4. Details about *pyr*/*ths* locus, related to Figure 5**

(A) Genome browser track visualized by IGV.[81] Publicly available ChIP-seq tracks for Zld in 120-150 min AEL embryos (GEO: GSE30757),[82] Dl in 2-4 h AEL embryos (GEO: GSE55306),[83] and Pol II in 2h10-2h40 AEL embryos (GEO: GSE295515)[57] were used for visualization. In the gene track, dark- and dim-colored isoforms represent active and inactive isoforms in early embryos, respectively. The region deleted in *Δths TSS* strain is highlighted.

(B) Relative expression levels of *pyr* and *ths* measured by qRT-PCR. Expression levels were normalized to *RpL32* and scaled to *yw* embryos. Bars represent the mean, and error bars represent the standard error of the mean. The p-values were calculated using a two-sided Welch's *t*-test. n = 13 for *yw* embryos; n = 13 for homozygous *Δths TSS* embryos. For *yw*, a part of the data is the same as that shown in Figures 5D and 5F.

Umemura *et al.*, Figure S5

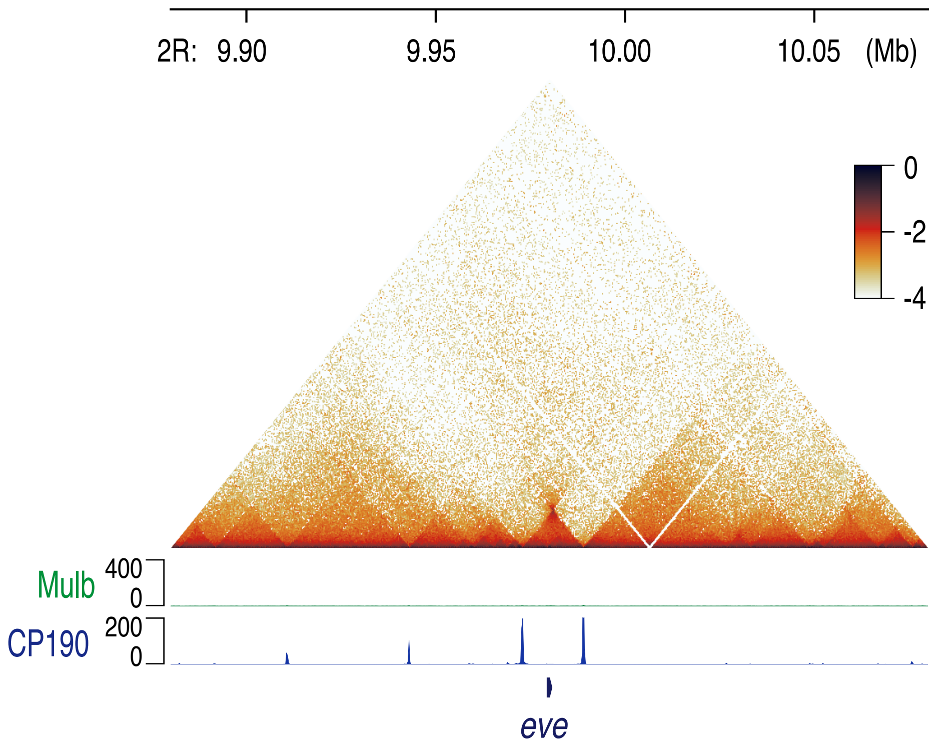

**Figure S5. Mulb-depleted *eve* locus, related to Figure 6**

Micro-C contact-frequency map is shown alongside ChIP-seq tracks. Coolbox toolkit[79] was used for visualization.

Umemura *et al.*, Figure S6

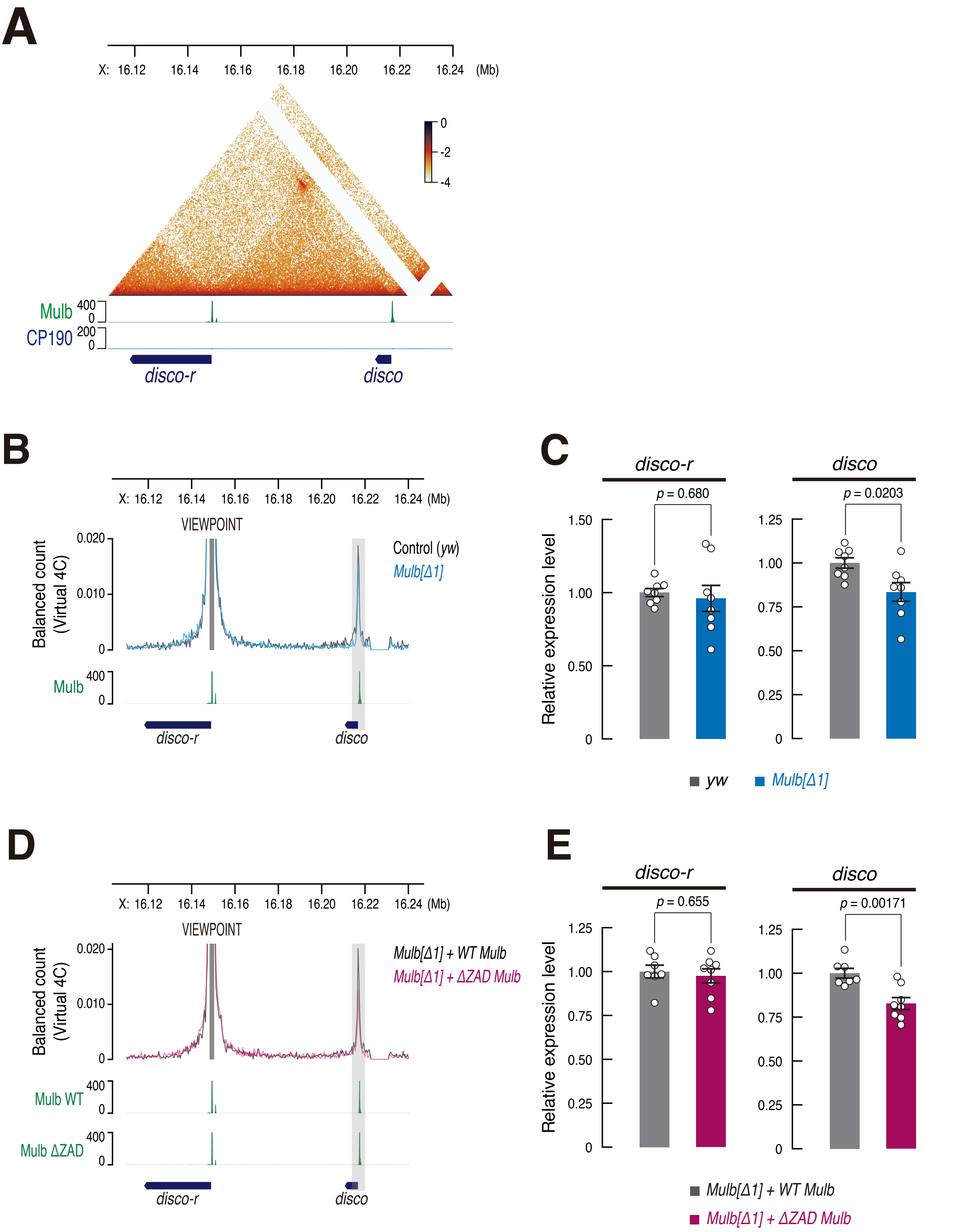

**Figure S6. Mulb fosters the formation of *disco*/*disco-r* promoter-promoter loop, related to Figure 6**

(A) Micro-C contact-frequency map is shown alongside ChIP-seq tracks. Micro-C experiment was performed using *yw* embryos. Coolbox toolkit[79] was used for visualization.

(B) Virtual 4C profiles in *yw* and homozygous *Mulb[Δ1]* embryos alongside ChIP-seq track of endogenous Mulb. For the viewpoint, a 2 kbp-scaled anchor of *disco*/*disco-r* promoter-promoter loop[3] (X:16148400-16150400) was used. Region around *disco* promoter is highlighted.

(C) Relative expression levels of *disco* and *disco-r* measured by qRT-PCR using 8-10 h AEL embryos. Measurements were performed with 8-10 h AEL embryos because *disco* and *disco-r* are not coactivated in nc14 embryos.[105] Expression levels were normalized to *RpL32* and scaled to *yw* embryos. Bars represent the mean, and error bars represent the standard error of the mean. The p-values were calculated using a two-sided Welch's *t*-test. n = 8 for *yw* embryos; n = 8 for homozygous *Mulb[Δ1]* embryos.

(D) Same as (B), but Virtual 4C profiles in homozygous rescue embryos (*Mulb[Δ1]* background) are shown alongside ChIP-seq tracks of WT and ΔZAD Mulb.

(E) Same as (C), but homozygous rescue embryos (*Mulb[Δ1]* background) were used. Expression levels were scaled to homozygous *WT Mulb* rescue embryos. n = 7 for homozygous *WT Mulb* rescue embryos; n = 8 for homozygous *ΔZAD Mulb* rescue embryos.

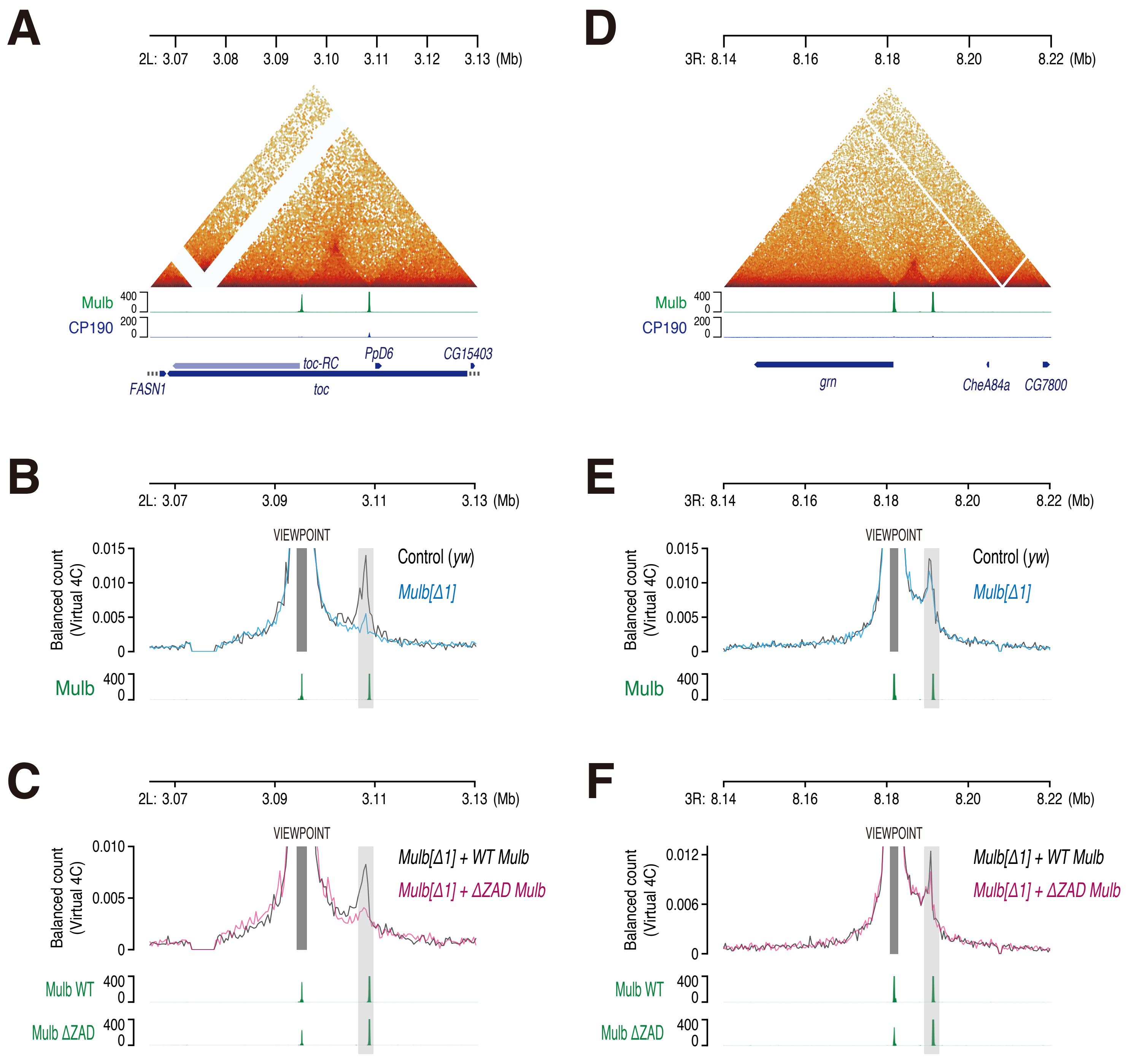
A
2L: 3.07
3.08
3.09
3.10
3.11
3.12
3.13 (Mb)
Mulb 400 0
CP190 200 0
toc-RC
PpD6
CG15403
FASN1
toc
D
3R: 8.14
8.16
8.18
8.20
8.22 (Mb)
Mulb 400 0
CP190 200 0
grn
CheA84a
CG7800
B
2L: 3.07
3.09
3.11
3.13 (Mb)
VIEWPOINT
Balanced count (Virtual 4C)
0.015
0.010
0.005
0
Control (yw)
Mulb[Δ1]
Mulb 400 0
E
3R: 8.14
8.16
8.18
8.20
8.22 (Mb)
VIEWPOINT
Balanced count (Virtual 4C)
0.015
0.010
0.005
0
Control (yw)
Mulb[Δ1]
Mulb 400 0
C
2L: 3.07
3.09
3.11
3.13 (Mb)
VIEWPOINT
Balanced count (Virtual 4C)
0.010
0.005
0
Mulb[Δ1] + WT Mulb
Mulb[Δ1] + ΔZAD Mulb
Mulb WT 400 0
Mulb ΔZAD 400 0
F
3R: 8.14
8.16
8.18
8.20
8.22 (Mb)
VIEWPOINT
Balanced count (Virtual 4C)
0.012
0.006
0
Mulb[Δ1] + WT Mulb
Mulb[Δ1] + ΔZAD Mulb
Mulb WT 400 0
Mulb ΔZAD 400 0

**Figure S7. Mulb regulates genome topology at non-paralogous loci, related to Figure 6**

(A) Micro-C contact-frequency map is shown alongside ChIP-seq tracks. Micro-C experiment was performed using *yw* embryos. Coolbox toolkit[79] was used for visualization. An isoform of *toc* with Mulb enrichment at the promoter region, *toc-RC*, is shown on the gene track.

(B) Virtual 4C profiles in *yw* and homozygous *Mulb[Δ1]* embryos alongside ChIP-seq track of Mulb. For the viewpoint, a 2 kbp-scaled anchor of the loop[3] (2L:3094400-3096400) was used. Region around the corresponding anchor is highlighted.

(C) Same as (B), but Virtual 4C profiles in homozygous rescue embryos (*Mulb[Δ1]* background) are shown alongside ChIP-seq tracks of WT and ΔZAD Mulb.

(D) Micro-C contact-frequency map is shown alongside ChIP-seq tracks. Micro-C experiment was performed using *yw* embryos. Coolbox toolkit[79] was used for visualization.

(E) Virtual 4C profiles in *yw* and homozygous *Mulb[Δ1]* embryos alongside ChIP-seq track of Mulb. For the viewpoint, a 2 kbp-scaled anchor of the loop[3] (3R:8180800-8182800) was used. Region around the corresponding anchor is highlighted.

(F) Same as (E), but Virtual 4C profiles in homozygous rescue embryos (*Mulb[Δ1]* background) are shown alongside ChIP-seq tracks of WT and ΔZAD Mulb.

#### Supplementary information

**Table S1. Sequences of smiFISH probes used in this study, related to Method Details.**

**Table S2. Sequences of DNA oligos used for gRNA cloning in this study, related to Method Details.**

**Table S3. Quality metrics for ChIP-seq libraries generated in this study, related to Method Details.**

**Table S4. Quality metrics for Micro-C libraries generated in this study, related to Method Details.**

**Video S1. Live-imaging of CG31365 in an early embryo, related to Figure 2**

Representative Airyscan time-lapse sequence showing the localization pattern of CG31365 from nc13 to nc14. Maximum projected images are shown. CG31365-GFP-3×FLAG and His2Av-emiRFP670 signals are shown in green and magenta, respectively.
