## Supplementary material for "The *Drosophila* ZAD zinc finger protein Mulberry shapes the organization of the regulatory genome in the early embryo": Table S1

Sequences of smiFISH probes used in this study, related to Method Details.

| *pyr* probes | |
|---|---|
| pyr_01 | 5′- CCTCCTAAGTTTCGAGCTGGACTCAGTGAGCGATCTACAACCGACGAA -3′ |
| pyr_02 | 5′- CCTCCTAAGTTTCGAGCTGGACTCAGTGTGTGTCTGGGACTTGGTATC -3′ |
| pyr_03 | 5′- CCTCCTAAGTTTCGAGCTGGACTCAGTGATTTTCCAAGGCATTTGGTG -3′ |
| pyr_04 | 5′- CCTCCTAAGTTTCGAGCTGGACTCAGTGGCTATCGATTGTGACTTGGA -3′ |
| pyr_05 | 5′- CCTCCTAAGTTTCGAGCTGGACTCAGTGATTAATCTGGCTCGAACGCA -3′ |
| pyr_06 | 5′- CCTCCTAAGTTTCGAGCTGGACTCAGTGAATGCACTTTTCACCTCTGG -3′ |
| pyr_07 | 5′- CCTCCTAAGTTTCGAGCTGGACTCAGTGAAATCAGTATCTGTAGCCGC -3′ |
| pyr_08 | 5′- CCTCCTAAGTTTCGAGCTGGACTCAGTGTTTTGGGTGTGCGATTTTTA -3′ |
| pyr_09 | 5′- CCTCCTAAGTTTCGAGCTGGACTCAGTGCACTGGGGCACTCGAAAATA -3′ |
| pyr_10 | 5′- CCTCCTAAGTTTCGAGCTGGACTCAGTGATCTTTTGTGTATTCTCGCG -3′ |
| pyr_11 | 5′- CCTCCTAAGTTTCGAGCTGGACTCAGTGCGCAACGACGACACGATATT -3′ |
| pyr_12 | 5′- CCTCCTAAGTTTCGAGCTGGACTCAGTGATTGCACTTTTGTCAACGGC -3′ |
| pyr_13 | 5′- CCTCCTAAGTTTCGAGCTGGACTCAGTGGCCTTGATTTGCGTATTTTA -3′ |
| pyr_14 | 5′- CCTCCTAAGTTTCGAGCTGGACTCAGTGCTTTATGCTATGTACATGCA -3′ |
| pyr_15 | 5′- CCTCCTAAGTTTCGAGCTGGACTCAGTGTTTTTGTATTTTTCCAACCG -3′ |
| pyr_16 | 5′- CCTCCTAAGTTTCGAGCTGGACTCAGTGTTGTTTTTCGTAACTCTTGC -3′ |
| pyr_17 | 5′- CCTCCTAAGTTTCGAGCTGGACTCAGTGCTTGCTTGGTTTTCAGTTTT -3′ |
| pyr_18 | 5′- CCTCCTAAGTTTCGAGCTGGACTCAGTGAAATGCAATCGAATCCGGGC -3′ |
| pyr_19 | 5′- CCTCCTAAGTTTCGAGCTGGACTCAGTGGCAGCGTCAACAAATTTACA -3′ |
| pyr_20 | 5′- CCTCCTAAGTTTCGAGCTGGACTCAGTGATCTGATACTTTAGGGCAGA -3′ |
| pyr_21 | 5′- CCTCCTAAGTTTCGAGCTGGACTCAGTGTAGATGTGTACGGCGAATCC -3′ |
| pyr_22 | 5′- CCTCCTAAGTTTCGAGCTGGACTCAGTGGTATAAACTTAGCGTATGCA -3′ |
| pyr_23 | 5′- CCTCCTAAGTTTCGAGCTGGACTCAGTGGTGTCCGAGCGGAAATTCAA -3′ |
| pyr_24 | 5′- CCTCCTAAGTTTCGAGCTGGACTCAGTGGTTTTAATTTTCTGCTGAGT -3′ |
| pyr_25 | 5′- CCTCCTAAGTTTCGAGCTGGACTCAGTGTCGTCGCAAGTCTATCATTT -3′ |
| pyr_26 | 5′- CCTCCTAAGTTTCGAGCTGGACTCAGTGAATCCAACATGGGCTTGTAC -3′ |
| pyr_27 | 5′- CCTCCTAAGTTTCGAGCTGGACTCAGTGCCCCCAAATTTAAGGGTATA -3′ |
| pyr_28 | 5′- CCTCCTAAGTTTCGAGCTGGACTCAGTGTATACTTTTGGCCGCATTGT -3′ |
| pyr_29 | 5′- CCTCCTAAGTTTCGAGCTGGACTCAGTGTATAAGTCCTGACATTGGGC -3′ |
| pyr_30 | 5′- CCTCCTAAGTTTCGAGCTGGACTCAGTGGCAATAGCAGCCACAAAACT -3′ |
| pyr_31 | 5′- CCTCCTAAGTTTCGAGCTGGACTCAGTGAAAACATTTTTCGCGGCGCT -3′ |
| pyr_32 | 5′- CCTCCTAAGTTTCGAGCTGGACTCAGTGTTCGCTGTGTTTGCTCAATG -3′ |
| pyr_33 | 5′- CCTCCTAAGTTTCGAGCTGGACTCAGTGCTTTGCCGTGCGAATTGATG -3′ |
| pyr_34 | 5′- CCTCCTAAGTTTCGAGCTGGACTCAGTGATATGTCCGCAGTATGTTCT -3′ |
| pyr_35 | 5′- CCTCCTAAGTTTCGAGCTGGACTCAGTGGACAGCTTCCATATTGAAGT -3′ |
| pyr_36 | 5′- CCTCCTAAGTTTCGAGCTGGACTCAGTGAATCCAGGCTGAACACATCG -3′ |
| pyr_37 | 5′- CCTCCTAAGTTTCGAGCTGGACTCAGTGCGCCGAATAGATGGTGATTT -3′ |
| pyr_38 | 5′- CCTCCTAAGTTTCGAGCTGGACTCAGTGGCCGTGATGGAAGCACAAAT -3′ |
| pyr_39 | 5′- CCTCCTAAGTTTCGAGCTGGACTCAGTGTAAAGTGGCACTCCTTTGTG -3′ |
| pyr_40 | 5′- CCTCCTAAGTTTCGAGCTGGACTCAGTGTGATTGCCCATGAAAATTCC -3′ |
| pyr_41 | 5′- CCTCCTAAGTTTCGAGCTGGACTCAGTGTGGATTGTAGGCGCTACTAA -3′ |
| pyr_42 | 5′- CCTCCTAAGTTTCGAGCTGGACTCAGTGGAAATTTCCCTTGAAACCCA -3′ |
| pyr_43 | 5′- CCTCCTAAGTTTCGAGCTGGACTCAGTGAATCATTCAAGCCAATGGGC -3′ |
| pyr_44 | 5′- CCTCCTAAGTTTCGAGCTGGACTCAGTGTTCTTCATCACTTTCGGTTT -3′ |
| pyr_45 | 5′- CCTCCTAAGTTTCGAGCTGGACTCAGTGATTTCTCCTCATTCAAAGGA -3′ |
| pyr_46 | 5′- CCTCCTAAGTTTCGAGCTGGACTCAGTGCAGCACTTCAGCCAAATGTA -3′ |
| pyr_47 | 5′- CCTCCTAAGTTTCGAGCTGGACTCAGTGTTTGGAACGCGTCCGTTTTG -3′ |
| pyr_48 | 5′- CCTCCTAAGTTTCGAGCTGGACTCAGTGTAATACTTGTTGGGCGACGC -3′ |

| *ths* probes | |
|---|---|
| ths_01 | 5′- CCTCCTAAGTTTCGAGCTGGACTCAGTGGGGCGATTGAATTATCGAGA -3′ |
| ths_02 | 5′- CCTCCTAAGTTTCGAGCTGGACTCAGTGCGAGACGTTGGACGTCTTGG -3′ |
| ths_03 | 5′- CCTCCTAAGTTTCGAGCTGGACTCAGTGAATGTCGGCTAAAGGCGTTG -3′ |
| ths_04 | 5′- CCTCCTAAGTTTCGAGCTGGACTCAGTGAAATCGCGTAATGCACGGGA -3′ |
| ths_05 | 5′- CCTCCTAAGTTTCGAGCTGGACTCAGTGATTTGCTAAATGTGGGGTGT -3′ |
| ths_06 | 5′- CCTCCTAAGTTTCGAGCTGGACTCAGTGACGTTTCTTTATTATTCGTT -3′ |
| ths_07 | 5′- CCTCCTAAGTTTCGAGCTGGACTCAGTGGTTTCAATACTTTATTGGGG -3′ |
| ths_08 | 5′- CCTCCTAAGTTTCGAGCTGGACTCAGTGGCTTTGCGCTTGCAATAATC -3′ |
| ths_09 | 5′- CCTCCTAAGTTTCGAGCTGGACTCAGTGACACACGCGATGCGATGTAC -3′ |
| ths_10 | 5′- CCTCCTAAGTTTCGAGCTGGACTCAGTGACACACATGTGAACACTCGC -3′ |
| ths_11 | 5′- CCTCCTAAGTTTCGAGCTGGACTCAGTGATTGCTGGCCGACAAATAGA -3′ |
| ths_12 | 5′- CCTCCTAAGTTTCGAGCTGGACTCAGTGGATTTATGTGCGTTCAGAGC -3′ |
| ths_13 | 5′- CCTCCTAAGTTTCGAGCTGGACTCAGTGCTATCCCCTCTATGTAATAT -3′ |
| ths_14 | 5′- CCTCCTAAGTTTCGAGCTGGACTCAGTGCGTGTAATCAGTTGACCGAT -3′ |
| ths_15 | 5′- CCTCCTAAGTTTCGAGCTGGACTCAGTGCTGACATTGGGCAGTTTCGG -3′ |
| ths_16 | 5′- CCTCCTAAGTTTCGAGCTGGACTCAGTGGTGATAACGCGCTGGCTAAC -3′ |
| ths_17 | 5′- CCTCCTAAGTTTCGAGCTGGACTCAGTGATTCAGAGGTGTCGTAGTTG -3′ |
| ths_18 | 5′- CCTCCTAAGTTTCGAGCTGGACTCAGTGATTTTAACCCATTTACTGCC -3′ |
| ths_19 | 5′- CCTCCTAAGTTTCGAGCTGGACTCAGTGCGACATTCATTCGGCGATTA -3′ |
| ths_20 | 5′- CCTCCTAAGTTTCGAGCTGGACTCAGTGACAGCAGTCTCTCTAACTGA -3′ |
| ths_21 | 5′- CCTCCTAAGTTTCGAGCTGGACTCAGTGAATGCACCTGAAATCACCAC -3′ |
| ths_22 | 5′- CCTCCTAAGTTTCGAGCTGGACTCAGTGGTGTGCACACTGTGACATTA -3′ |
| ths_23 | 5′- CCTCCTAAGTTTCGAGCTGGACTCAGTGCAGCTATGTGTATGGCTTTG -3′ |
| ths_24 | 5′- CCTCCTAAGTTTCGAGCTGGACTCAGTGCTGTGAAGTGTCTGTCACTG -3′ |
| ths_25 | 5′- CCTCCTAAGTTTCGAGCTGGACTCAGTGGCCGACTATCGTTATATTTT -3′ |
| ths_26 | 5′- CCTCCTAAGTTTCGAGCTGGACTCAGTGTCGTTGTTCAGATAATCCGG -3′ |
| ths_27 | 5′- CCTCCTAAGTTTCGAGCTGGACTCAGTGTGCGTACAGTGCTATTTTGA -3′ |
| ths_28 | 5′- CCTCCTAAGTTTCGAGCTGGACTCAGTGGCCAACTAATCTCCAATTGT -3′ |
| ths_29 | 5′- CCTCCTAAGTTTCGAGCTGGACTCAGTGGTCTCGTTGAAATAGCAGGT -3′ |
| ths_30 | 5′- CCTCCTAAGTTTCGAGCTGGACTCAGTGCACGAAATATCCGTGCACGA -3′ |
| ths_31 | 5′- CCTCCTAAGTTTCGAGCTGGACTCAGTGGCAGATCGACAACGGAACGG -3′ |
| ths_32 | 5′- CCTCCTAAGTTTCGAGCTGGACTCAGTGCATCGTTGACGCTCTTCTTG -3′ |
| ths_33 | 5′- CCTCCTAAGTTTCGAGCTGGACTCAGTGATCTTGTTGAACATGTAGCA -3′ |
| ths_34 | 5′- CCTCCTAAGTTTCGAGCTGGACTCAGTGATGGCGGAAAAACTGCTCGG -3′ |
| ths_35 | 5′- CCTCCTAAGTTTCGAGCTGGACTCAGTGCACTGTTGCCATTGTTATGG -3′ |
| ths_36 | 5′- CCTCCTAAGTTTCGAGCTGGACTCAGTGTTGTTTTTGTTGTTGCGCGG -3′ |
| ths_37 | 5′- CCTCCTAAGTTTCGAGCTGGACTCAGTGTTGGTTATTCGATTTGTGGC -3′ |
| ths_38 | 5′- CCTCCTAAGTTTCGAGCTGGACTCAGTGCTCCATTTCGGTGATGTAGG -3′ |
| ths_39 | 5′- CCTCCTAAGTTTCGAGCTGGACTCAGTGTACTGCATCAACTCGGTGTC -3′ |
| ths_40 | 5′- CCTCCTAAGTTTCGAGCTGGACTCAGTGGAATTGTCCAGGTGATCTTC -3′ |
| ths_41 | 5′- CCTCCTAAGTTTCGAGCTGGACTCAGTGTGTTGTGGCTTCACTTCGTT -3′ |
| ths_42 | 5′- CCTCCTAAGTTTCGAGCTGGACTCAGTGCACTGTCCACAGTAACTTGC -3′ |
| ths_43 | 5′- CCTCCTAAGTTTCGAGCTGGACTCAGTGCAATGTTGCTAGTGTGCTGC -3′ |
| ths_44 | 5′- CCTCCTAAGTTTCGAGCTGGACTCAGTGAGTGACGGTGCAACATGTTG -3′ |
| ths_45 | 5′- CCTCCTAAGTTTCGAGCTGGACTCAGTGTGTTGATGTTCTTGTCCACG -3′ |
| ths_46 | 5′- CCTCCTAAGTTTCGAGCTGGACTCAGTGGTTAGTGTGTTGTTCAGGTT -3′ |
| ths_47 | 5′- CCTCCTAAGTTTCGAGCTGGACTCAGTGTGATGTTGCCGTGGGAGAAG -3′ |
| ths_48 | 5′- CCTCCTAAGTTTCGAGCTGGACTCAGTGTTGCGGACGTGATGCTGGAT -3′ |

| *eve* probes | |
|---|---|
| eve_01 | 5′- CCTCCTAAGTTTCGAGCTGGACTCAGTGAGGCAGTTAGTTGTTGACTG -3′ |
| eve_02 | 5′- CCTCCTAAGTTTCGAGCTGGACTCAGTGAGTTGGCTTATTCAGAGGAT -3′ |
| eve_03 | 5′- CCTCCTAAGTTTCGAGCTGGACTCAGTGTTGGTATGCGTCTTGTGATT -3′ |
| eve_04 | 5′- CCTCCTAAGTTTCGAGCTGGACTCAGTGCATGTTGTAGGTTCGGTATC -3′ |
| eve_05 | 5′- CCTCCTAAGTTTCGAGCTGGACTCAGTGCAAGAGGTCCACAACCAGGG -3′ |
| eve_06 | 5′- CCTCCTAAGTTTCGAGCTGGACTCAGTGTTCAAGGAGTTATCCGGACT -3′ |
| eve_07 | 5′- CCTCCTAAGTTTCGAGCTGGACTCAGTGTGTAGAACTCCTTCTCCAAG -3′ |
| eve_08 | 5′- CCTCCTAAGTTTCGAGCTGGACTCAGTGGGACGGGACACGTAGTTCTC -3′ |
| eve_09 | 5′- CCTCCTAAGTTTCGAGCTGGACTCAGTGTTCTGGAACCACACCTTGAT -3′ |
| eve_10 | 5′- CCTCCTAAGTTTCGAGCTGGACTCAGTGTCTGACGCTTGTCCTTCATG -3′ |
| eve_11 | 5′- CCTCCTAAGTTTCGAGCTGGACTCAGTGAAGGCGGGATCGGAGTAGAC -3′ |
| eve_12 | 5′- CCTCCTAAGTTTCGAGCTGGACTCAGTGATCTGCATTGTGGGCATCTG -3′ |
| eve_13 | 5′- CCTCCTAAGTTTCGAGCTGGACTCAGTGTGTAGCGGTACTGTCCGTAG -3′ |
| eve_14 | 5′- CCTCCTAAGTTTCGAGCTGGACTCAGTGTATGAGGACCAGCGGGATGT -3′ |
| eve_15 | 5′- CCTCCTAAGTTTCGAGCTGGACTCAGTGCCCATCATGTGCGGATGATG -3′ |
| eve_16 | 5′- CCTCCTAAGTTTCGAGCTGGACTCAGTGGATGACGACGACTGCAGATC -3′ |
| eve_17 | 5′- CCTCCTAAGTTTCGAGCTGGACTCAGTGAGAGCGACAGCGTGGAGGAG -3′ |
| eve_18 | 5′- CCTCCTAAGTTTCGAGCTGGACTCAGTGTCGAACACCTTGGCGTGATC -3′ |
| eve_19 | 5′- CCTCCTAAGTTTCGAGCTGGACTCAGTGCGGATTGAGCCACTGGACTG -3′ |
| eve_20 | 5′- CCTCCTAAGTTTCGAGCTGGACTCAGTGAGCGGGAGCAGGAACTGATG -3′ |
| eve_21 | 5′- CCTCCTAAGTTTCGAGCTGGACTCAGTGCGCAATCACAGTTGTCGTCG -3′ |
| eve_22 | 5′- CCTCCTAAGTTTCGAGCTGGACTCAGTGCTTGAAGAGCTTCGGCTTGG -3′ |
| eve_23 | 5′- CCTCCTAAGTTTCGAGCTGGACTCAGTGCTTACGCCTCAGTCTTGTAG -3′ |
| eve_24 | 5′- CCTCCTAAGTTTCGAGCTGGACTCAGTGGAGAGAGTGTGTGTGGATCG -3′ |
| eve_25 | 5′- CCTCCTAAGTTTCGAGCTGGACTCAGTGTACAATCTTTTGGGGGAGCA -3′ |
| eve_26 | 5′- CCTCCTAAGTTTCGAGCTGGACTCAGTGGAGGCTGACTAAGACTAGTT -3′ |
| eve_27 | 5′- CCTCCTAAGTTTCGAGCTGGACTCAGTGTCTGTACAATCTTCGGGAAT -3′ |
| eve_28 | 5′- CCTCCTAAGTTTCGAGCTGGACTCAGTGTGACTACAATTAGCTACTCT -3′ |
| eve_29 | 5′- CCTCCTAAGTTTCGAGCTGGACTCAGTGCTTAATTTGATTTTGCGCCT -3′ |
| eve_30 | 5′- CCTCCTAAGTTTCGAGCTGGACTCAGTGGGTGAAGGCGGTGCGATAG -3′ |
| eve_31 | 5′- CCTCCTAAGTTTCGAGCTGGACTCAGTGTAGGGCGGATAGGGCATGC -3′ |
| eve_32 | 5′- CCTCCTAAGTTTCGAGCTGGACTCAGTGGAGTACGACGATCCCGTTG -3′ |
| eve_33 | 5′- CCTCCTAAGTTTCGAGCTGGACTCAGTGAGGGCAGAGCGCCCAAAAG -3′ |
| eve_34 | 5′- CCTCCTAAGTTTCGAGCTGGACTCAGTGCAGTCCGGTATAGCAGGTG -3′ |
| eve_35 | 5′- CCTCCTAAGTTTCGAGCTGGACTCAGTGTGGCACTGGGCATCAGGAG -3′ |

| MS2 probes | |
|---|---|
| MS2_01 | 5′- CCTCCTAAGTTTCGAGCTGGACTCAGTGTCGTGCTTTCTTGGCAATAA -3′ |
| MS2_02 | 5′- CCTCCTAAGTTTCGAGCTGGACTCAGTGCGTTTGAAGATTCGACCTGG -3′ |
| MS2_03 | 5′- CCTCCTAAGTTTCGAGCTGGACTCAGTGAATACTGGAGCGACGCGTGA -3′ |
| MS2_04 | 5′- CCTCCTAAGTTTCGAGCTGGACTCAGTGACCGTAGGATCTGATGAACC -3′ |
