## Supplementary material for "The *Drosophila* ZAD zinc finger protein Mulberry shapes the organization of the regulatory genome in the early embryo": Table S2

Sequences of DNA oligos used for gRNA cloning in this study, related to Method Details.

| gRNA | Sense oligo | Antisense oligo |
|---|---|---|
| CG31365[Δ1]_1 | 5′- GTCGAAATCAACGTATTTACTCGG -3′ | 5′- AAACCCGAGTAAATACGTTGATTT -3′ |
| CG31365[Δ1]_2 | 5′- GTCGAAACATGGTGGTCCAATCTG -3′ | 5′- AAACCAGATTGGACCACCATGTTT -3′ |
| CG31365-GFP-3xFLAG | 5′- GTCGACGAACGAGTTTCAAGCTG -3′ | 5′- AAACCAGCTTGAAACTCGTTCGT -3′ |
| ΔMulbSite_1 | 5′- GTCGGGCTTCAGTCTCAAGTGGTT -3′ | 5′- AAACAACCACTTGAGACTGAAGCC -3′ |
| ΔMulbSite_2 | 5′- GTCGTAACTGTTCTCGCATATCAT -3′ | 5′- AAACATGATATGCGAGAACAGTTA -3′ |
| ΔEnhancer_1 | 5′- GTCGAAAGTACCTATGAATGGTAA -3′ | 5′- AAACTTACCATTCATAGGTACTTT -3′ |
| ΔEnhancer_2 | 5′- GTCGGCCGGAATATATAGGGGAGC -3′ | 5′- AAACGCTCCCCTATATATTCCGGC -3′ |
| ΔthsTSS_1 | 5′- GTCGGGTGGATGATGTCCACGAGT -3′ | 5′- AAACACTCGTGGACATCATCCACC -3′ |
| ΔthsTSS_2 | 5′- GTCGGTCGAAAACCCCGCTACCCT -3′ | 5′- AAACAGGGTAGCGGGGTTTTCGAC -3′ |
