## Supplementary material for "The *Drosophila* ZAD zinc finger protein Mulberry shapes the organization of the regulatory genome in the early embryo": Table S3

Quality metrics for ChIP-seq libraries generated in this study, related to Method Details.

| Library | Total reads number | Reads number after filtering (fastp) | Overall alignment rate (bowtie2) |
|---|---:|---:|---:|
| Mulb_FLAG-ChIP_rep1_ChIP | 42,039,438 | 42,039,078 | 95.83% |
| Mulb_FLAG-ChIP_rep1_input | 12,001,620 | 12,001,596 | 97.62% |
| Mulb_FLAG-ChIP_rep2_ChIP | 22,459,572 | 22,459,428 | 95.30% |
| Mulb_FLAG-ChIP_rep2_input | 9,507,870 | 9,507,858 | 97.70% |
| CP190_FLAG-ChIP_rep1_ChIP | 66,241,008 | 66,240,746 | 97.06% |
| CP190_FLAG-ChIP_rep1_input | 5,933,754 | 5,933,746 | 97.97% |
| CP190_FLAG-ChIP_rep2_ChIP | 46,822,298 | 46,822,036 | 95.19% |
| CP190_FLAG-ChIP_rep2_input | 14,257,012 | 14,256,996 | 98.22% |
| WTMulbinMulbKO_FLAG-ChIP_rep1_ChIP | 34,053,004 | 34,052,092 | 95.56% |
| WTMulbinMulbKO_FLAG-ChIP_rep1_input | 21,256,602 | 21,256,530 | 97.08% |
| WTMulbinMulbKO_FLAG-ChIP_rep2_ChIP | 46,232,018 | 46,231,622 | 96.16% |
| WTMulbinMulbKO_FLAG-ChIP_rep2_input | 18,843,938 | 18,843,916 | 97.18% |
| delZADMulbinMulbKO_FLAG-ChIP_rep1_ChIP | 20,810,584 | 20,810,230 | 94.58% |
| delZADMulbinMulbKO_FLAG-ChIP_rep1_input | 18,386,296 | 18,386,260 | 96.92% |
| delZADMulbinMulbKO_FLAG-ChIP_rep2_ChIP | 21,257,960 | 21,257,476 | 94.97% |
| delZADMulbinMulbKO_FLAG-ChIP_rep2_input | 19,001,578 | 19,001,554 | 97.24% |
| yw_FLAG-ChIP_rep1_ChIP | 7,246,230 | 7,246,028 | 84.94% |
| yw_FLAG-ChIP_rep1_input | 4,855,180 | 4,855,168 | 97.01% |
| yw_FLAG-ChIP_rep2_ChIP | 10,488,386 | 10,488,216 | 92.58% |
| yw_FLAG-ChIP_rep2_input | 9,957,782 | 9,957,762 | 96.60% |
