## Supplementary material for "The *Drosophila* ZAD zinc finger protein Mulberry shapes the organization of the regulatory genome in the early embryo": Table S4

Quality metrics for Micro-C libraries generated in this study, related to Method Details.

| Library | Total read pairs (fastp) | Read pairs number after filtering (fastp) | Total mapped pairs (pairtools) | Total mapped pairs without duplicates (pairtools) | Total *cis* pairs (pairtools) | Total *trans* pairs (pairtools) | Total mapped pairs / Read pairs number after filtering | Total cis pairs / Total mapped pairs without duplicates |
|---|---:|---:|---:|---:|---:|---:|---:|---:|
| yw_Micro-C_rep1 | 489,783,035 | 489,782,053 | 322,330,453 | 183,811,328 | 175,218,222 | 8,593,106 | 65.81% | 95.33% |
| yw_Micro-C_rep2 | 483,564,202 | 483,563,358 | 323,256,209 | 142,390,386 | 133,691,823 | 8,698,563 | 66.85% | 93.89% |
| MulbKO_Micro-C_rep1 | 494,858,408 | 494,858,207 | 334,492,904 | 294,008,833 | 281,144,481 | 12,864,352 | 67.59% | 95.62% |
| MulbKO_Micro-C_rep2 | 504,934,365 | 504,933,934 | 336,123,680 | 295,510,045 | 283,749,738 | 11,760,307 | 66.57% | 96.02% |
| WTMulbinMulbKO_Micro-C_rep1 | 505,141,696 | 505,125,669 | 305,033,204 | 162,423,194 | 117,602,752 | 44,820,442 | 60.39% | 72.41% |
| WTMulbinMulbKO_Micro-C_rep2 | 523,961,467 | 523,960,965 | 353,296,195 | 266,561,994 | 251,787,120 | 14,774,874 | 67.43% | 94.46% |
| delZADMulbinMulbKO_Micro-C_rep1 | 530,584,807 | 530,580,276 | 330,532,193 | 140,267,341 | 108,622,511 | 31,644,830 | 62.30% | 77.44% |
| delZADMulbinMulbKO_Micro-C_rep2 | 529,827,693 | 529,826,831 | 346,459,082 | 198,603,346 | 188,585,323 | 10,018,023 | 65.39% | 94.96% |
| delMulbSite_Micro-C_rep1 | 461,225,996 | 461,225,147 | 306,295,526 | 179,713,044 | 170,913,865 | 8,799,179 | 66.41% | 95.10% |
| delMulbSite_Micro-C_rep2 | 467,625,867 | 467,625,304 | 309,277,533 | 131,257,861 | 123,117,313 | 8,140,548 | 66.14% | 93.80% |
| delEnhancer_Micro-C_rep1 | 463,845,352 | 463,007,531 | 298,361,409 | 163,119,373 | 155,293,483 | 7,825,890 | 64.44% | 95.20% |
| delEnhancer_Micro-C_rep2 | 476,301,528 | 476,300,026 | 296,737,665 | 125,928,859 | 117,712,463 | 8,216,396 | 62.30% | 93.48% |
